# Spatiotemporal resonance in mouse primary visual cortex

**DOI:** 10.1101/2023.07.31.551212

**Authors:** Rasa Gulbinaite, Mojtaba Nazari, Michael E. Rule, Edgar J. Bermudez-Contreras, Michael X Cohen, Majid H. Mohajerani, J. Alexander Heimel

**Author notes:** These authors contributed equally (Majid H. Mohajerani, J. Alexander Heimel). The co-last authorship order was determined by a coin flip.

## Abstract

Human primary visual cortex (V1) responds more strongly, or resonates, when exposed to ∼10, ∼15-20, ∼40-50 Hz rhythmic flickering light. Full-field flicker also evokes perception of hallucinatory geometric patterns, which mathematical models explain as standing-wave formations emerging from periodic forcing at resonant frequencies of the simulated neural network. However, empirical evidence for such flicker-induced standing waves in the visual cortex was missing. We recorded cortical responses to flicker in awake mice using high spatial resolution widefield imaging in combination with high temporal resolution glutamate-sensing fluorescent reporter (iGluSnFR). The temporal frequency tuning curves in the mouse V1 were similar to those observed in humans, showing a banded structure with multiple resonance peaks (8 Hz, 15 Hz, 33 Hz). Spatially, all flicker frequencies evoked responses in V1 corresponding to retinotopic stimulus location, but some evoked additional peaks. These flicker-induced cortical patterns displayed standing wave characteristics and matched linear wave equation solutions in an area restricted to the visual cortex. Taken together, the interaction of periodic traveling waves with cortical area boundaries leads to spatiotemporal activity patterns that may affect perception.

## INTRODUCTION

Sensory neurons of many species – from fruit flies, rodents, salamanders to humans – stimulated with rhythmic sounds, lights, or touch closely follow the rhythm. This manifests as peaks in power spectra corresponding to the stimulus frequency and its harmonics in electroencephalogram (EEG), local field potentials (LFPs), and single-unit recordings^1–6^. In addition to the frequency-following neural response, a few notable response nonlinearities exist. In human EEG, stronger amplitude responses, or *resonance*, over occipitoparietal areas have been consistently reported for ∼10, 15–20, and 40–50 Hz rhythmic visual stimulation (flicker)^5,7,8^. Relatedly, the phase of steady-state visual evoked potentials (ssVEPs) at resonance frequencies is more stable^8,9^, suggesting entrainment of endogenous brain oscillators operating in the same frequency range^10^. Entrainment of intrinsic gamma to rhythmic 40 Hz stimuli was also put forward as a mechanism for reduced plaque formation in Alzheimer’s disease mouse model^11^.

While multiple resonance peaks are observed in the temporal frequency tuning curves in the human^5,7,8^ and cat visual cortex^4^, multiunit and spiking activity in the mouse visual cortex seems to show only a single (∼6 Hz) peak^12,13^. Similarly, two-photon calcium imaging studies indicate that spatiotemporal tuning in mouse V1 is restricted to low temporal frequencies^14^. Variable temporal resolution of methods and limited stimulation frequencies used in mouse studies may have concealed resonance frequencies and hindered direct comparisons of temporal frequency tuning across species. However, such comparisons are relevant for validating the mouse as a model for understanding neurophysiological mechanisms of ssVEP generation in humans and its relation to endogenous brain oscillations^15,16^, as well as therapeutic effects of non-invasive sensory stimulation on aberrant brain oscillations^17^.

In addition to temporal resonance, the spatial distribution of responses varies with flicker frequency. In fMRI studies, this is linked to the spatial organization of temporal frequency preference in the visual cortex^18,19^, whereas EEG studies attribute spatiotemporal response variability to varying wavelength traveling and standing waves^20,21^. Full-field luminance flicker also induces geometric visual hallucinations. In mathematical models^22^, these hallucinatory patterns are explained as standing waves occurring at resonance frequencies of the neural network. External rhythmic stimulation disrupts the network’s excitation-inhibition balance, causing intrinsic network oscillations to self-organize into standing waves. Although this theoretical pattern formation mechanism is based on anatomical knowledge (lateral connections, retinotopy and cortical magnification), the emergence of spatiotemporal wave phenomena on the cortical surface in response to flicker stimulation has not been demonstrated experimentally.

Although both standing and traveling waves have been reported in flicker stimulation paradigms in humans^20,21^, the limited spatial resolution of EEG and source mixing in scalp projections makes it difficult to unambiguously address the spatial organization of responses to different frequency flicker and its relation to wave phenomena^23,25^. To circumvent these limitations, we made use of the smooth-surfaced mouse cortex. We measured spatiotemporal propagation of responses to flicker at the mesoscale using a fast genetically encoded glutamate sensor iGluSnFR expressed in excitatory neurons across all cortical layers^24,26,27^(Figure 1A-B). To directly compare temporal frequency tuning in human and mouse V1, we employed a protocol similar to that used in our previous human study^8^.

**Figure 1.**
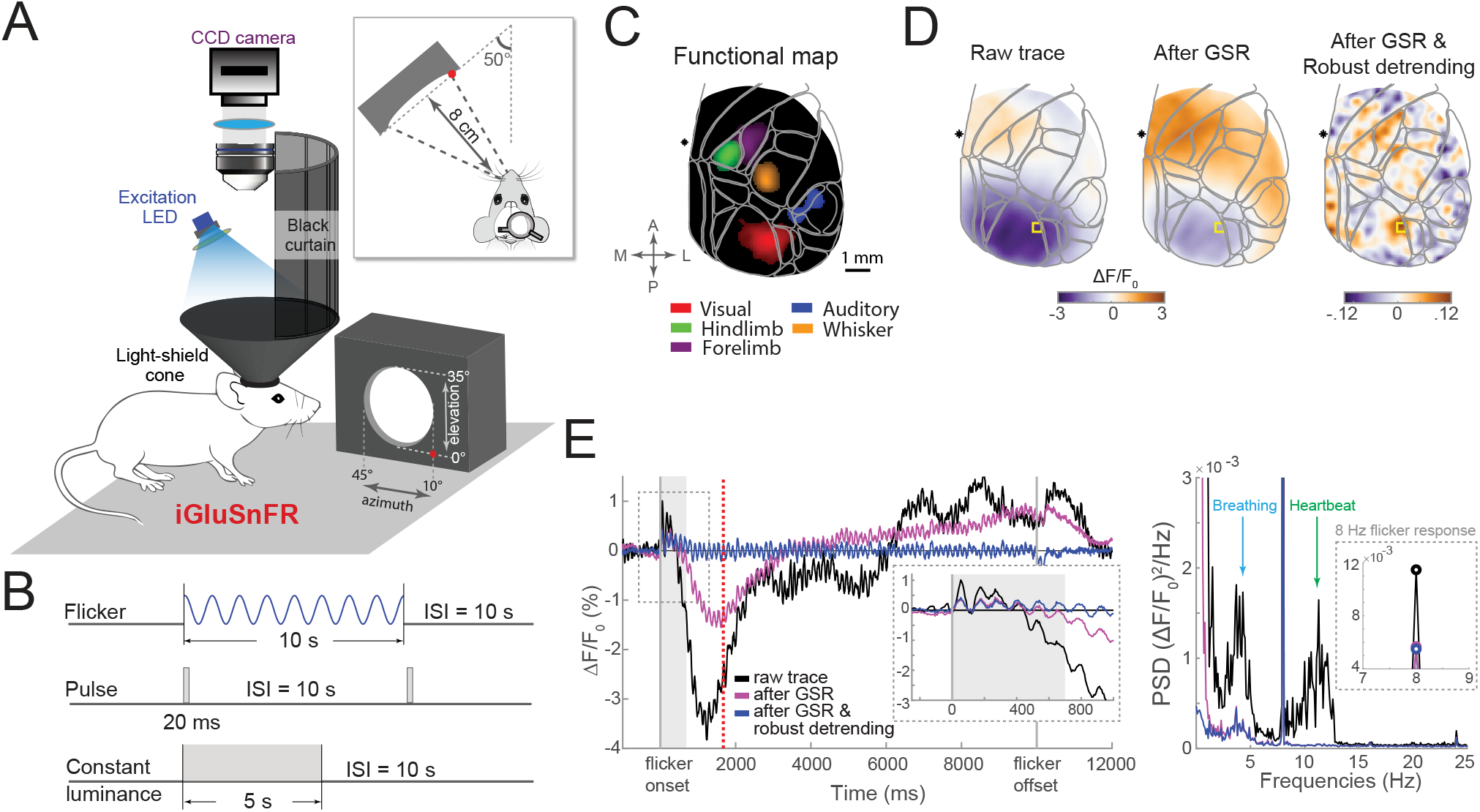
iGluSnFR fluorescence imaging and experimental design. (A) Schematic imaging setup and visual stimulation device. (B) Visual stimulation protocols. (C) Functional cortical map showing peak evoked activity from different sensory stimuli (single-animal data; thresholded at 5 SD relative to the baseline), co-registered to the Allen Mouse Brain Common Coordinate Framework (CCF; light gray outlines). (D) Cortical activity maps at the timepoint marked by the red dashed line in panel E. Note V1 suppression in baseline-corrected (raw trace) and GSR-corrected signals, and the expected V1 activation after robust detrending. (E) Left: Raw trace of trial-average response to 8 Hz flicker (black line), after global signal regression (GSR; magenta line) and robust detrending (blue line) corrections. Inset: zoomed-in time courses. Grey area (0-700 ms post-flicker onset) marks the time window where robust detrending was not applied to avoid removal of a neural signal. Right: Power spectra of the signals on the left show that GSR removed heartbeat and attenuated breathing-related artifacts, while robust detrending removed slow oscillations related to neurovascular coupling (a steep slope at low frequencies). See also Figures S1,S2.

## RESULTS

### Multiple temporal resonance frequencies in mouse and human V1

As expected based on the retinotopic organization of the visual cortex, left-hemifield flickering stimulus elicited retinotopically localized response over the right V1 (Figure 1C,2A). No activity in V1 was detected when iGluSnFR was not directly excited by the blue LED light (Figure S1E). To separate stimulus-related fluorescence changes (steady-state visual evoked responses, ssVERs) from respiratory and hemodynamic artifacts (heartbeat and neurovascular coupling) caused by an overlap between the excitation spectrum of iGluSnFR and the absorption spectrum of hemoglobin (Figure 1D,E)^24,28^, we applied global signal regression (GSR; STAR Methods)^24^. GSR removed contamination from the 8-14 Hz heartbeat signal, attenuated breathing-related artifacts, and improved overall signal-to-noise ratio (Figure 1E,S2C). However, it only partially attenuated the slow BOLD-like signal (<1 Hz), which we corrected with robust detrending (Figure 1E)^29,30^. After these corrections, fluorescence time courses closely followed sine-wave modulated stimulus luminance changes.

Power spectra from V1 pixels most responsive to the flicker (V1 ROI) showed a strong peak at the stimulus and harmonically related frequencies (Figures 2A,B). Peaks at half the stimulation frequency (subharmonics) were also present in response to 36–52 Hz flicker (Figure 2B), and likely reflect retinal ganglion cell firing to every other cycle of luminance increase (period doubling)^2,31^. Harmonic and subharmonic responses were distinguishable only after 1/*f* power-law scaling effect (smaller amplitude brain activity at higher frequencies) was accounted for by expressing the entire power spectra in signal-to-noise ratio (SNR) units (STAR Methods, Figure S1C,D). Conversion to SNR units facilitated comparison of responses across flicker frequencies and animals^16^. Statistically significant ssVERs in V1 were elicited up to 56 Hz flicker for 1*f* and 66 Hz for 2*f* responses (Figures 2C, S2E,F). Statistical significance of responses up to 56 Hz flicker was also confirmed at the cortical map level (Figure 2E,F). Thus, temporal resolution of iGluSnFR observed here in chronic *in vivo* recordings (up to 66 Hz) matched those reported *in vitro* in hippocampal slice preparations^27^ (⊠_1/2_ = 15⊠11 ms; 1/15·1000 = 67 Hz), and allowed us to track responses to gamma-band flicker stimuli. Higher-order visual area LM (lateromedial) responded up to 30 Hz flicker and PM (posteromedial) up to 18 Hz (Figure S2G), consistent with their respective preferences for fast- and slow-varying stimuli^14^. Additionally, slow (<11 Hz) flicker elicited small but statistically significant responses in non-visual areas (Figure 2E,F).

**Figure 2.**
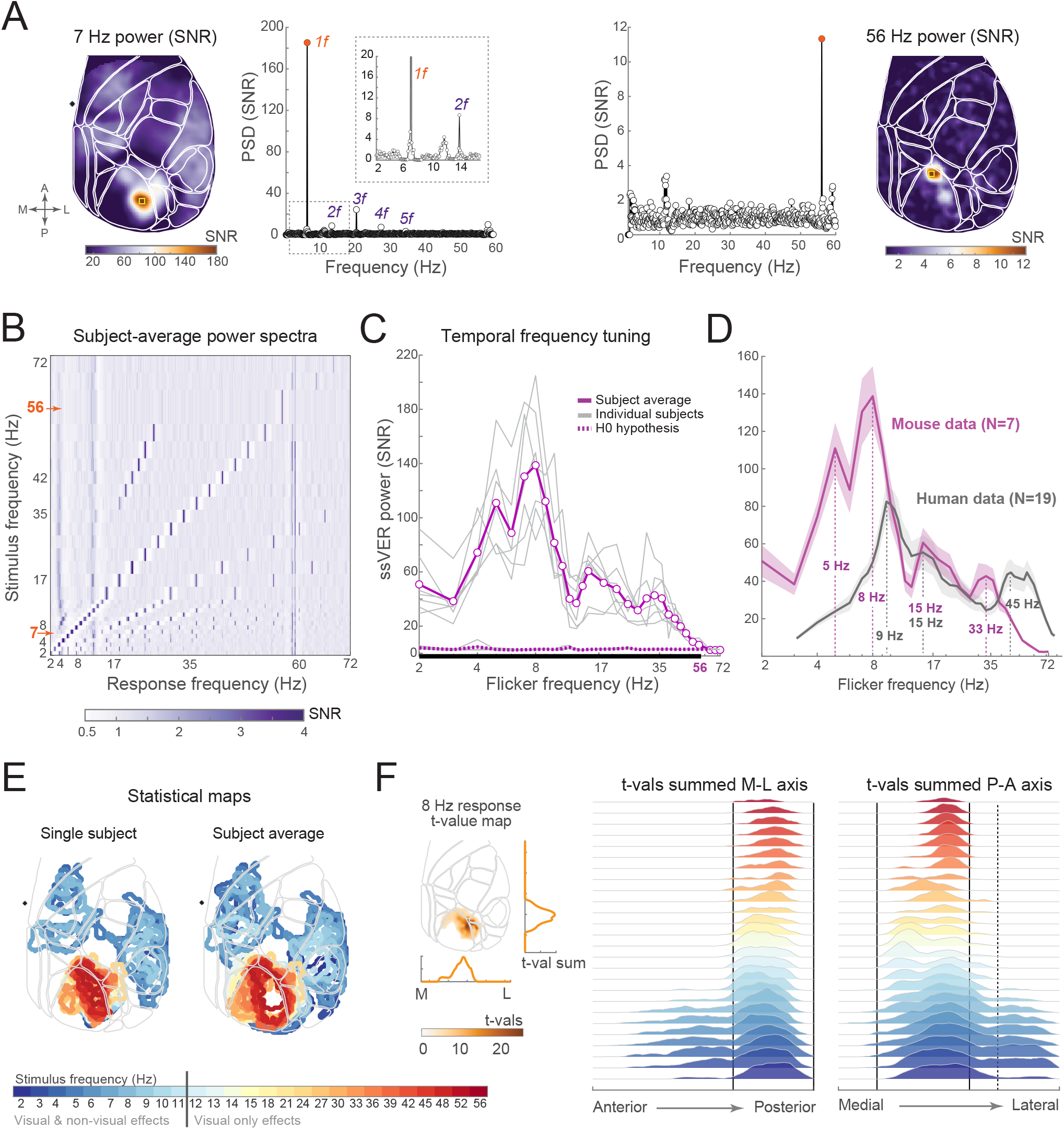
Temporal frequency tuning in the mouse visual cortex. (A) Topographies and SNR power spectra from V1 ROI (yellow square) in response to 7 Hz (left) and 56 Hz (right) flicker (single-animal data). Inset: zoomed-in power spectrum around the flicker frequency. (B) Subject-average power spectra from V1 ROI (x-axis) plotted as a function of flicker frequency (y-axis). Note the frequency-following response along the diagonal, harmonic (2*f*, 3*f*, etc.) and subharmonic *f/2* responses, as well as increased power at: (1) 3–5 Hz (see the main text); (2) 11–13 Hz (residual heartbeat artifact); (3) ∼57 Hz (CCD camera noise). (C) ssVER power across all 30 tested flicker frequencies: Single-animal (gray lines) and group-average data (N=7, magenta line). Dotted magenta line represents ssVER power on 95^th^ percentile of trials during which neither flicker frequency nor harmonically related frequencies were presented (H0 hypothesis). Thick black line marks significant ssVERs (up to 56 Hz flicker). (D) Comparison of temporal frequency tuning curves in mice (N=7) and humans (N=19; Gulbinaite et al., 2019). Shaded areas represent SEM. (E) Spatial extent of statistically significant responses as a function of flicker frequency. Statistical significance was determined using bootstrap-t procedure (see STAR Methods). Responses to flicker frequencies <11 Hz were present beyond visual areas (auditory and somatosensory). (F) Data from (E) projected onto medial-lateral (left) and anterior-posterior (right) axes by summing the observed t-values. Color-coded by flicker frequency. Inset illustrates the summing procedure. Vertical lines indicate boundaries of V1 (solid line) and lateral visual areas (dashed line). See also Figures S2,S3.

Although response amplitude decreased as flicker frequency increased, ssVER amplitude decrease was not monotonic. The grand-average temporal frequency tuning curves, constructed by plotting response power at each flicker frequency, contained resonance peaks in response to theta (M = 4.86 Hz, SD = 0.38 and M = 7.57 Hz; SD = 0.53 Hz), beta (M = 16.71 Hz, SD = 2.93 Hz), and gamma-band flicker (M = 33.86 Hz, SD = 3.34 Hz; Figure 2C). Gamma-band resonance peaks were most variable across animals (peak range 30-39 Hz). In contrast, temporal frequency tuning curves derived from raw rather than SNR power spectra exhibited only a single peak at 4–5 Hz (Figure S2B). We recomputed temporal frequency tuning curves without GSR to evaluate the impact of heart-beat artifacts on resonance peaks. As expected, responses to theta-band flicker were smaller and the resonance peak at ∼8 Hz was nearly absent because a strong heartbeat artifact affected SNR computation (Figure S2C,D). To further control for heart-beat artifact effects on resonance peaks, we computed temporal frequency tuning curves from visual and non-visual ROIs placed over the blood vessels (Figure S2H). Only the visual ROI contained resonance peaks (Figure S2I).

In addition to three resonance peaks reported in humans^5,8^ (Figure 2D), a fourth ∼5 Hz peak in the theta band was present in mice (Figure 2C,D). Although Hartigan’s dip test for multimodality suggested a single rather than a double peak in the theta band (∼5 Hz and ∼8 Hz peaks not different; p = 0.091), endogenous 3-5 Hz oscillations are prominent in mouse V1 LFP and single-cell recordings, with the strongest power in layer 2/3^32,33^. The 3-5 Hz rhythm was also present in iGluSnFR signal (Figures 1E,S3). Topographically, the 3-5 Hz rhythm was not constrained to visual areas (Figure S3A), was stronger during low- (< 20 Hz) vs. high-frequency flicker trials (Figure S3C), and more prominent during flicker vs. inter-trial interval (Figures S3B,D). We speculate that the 3–5 Hz rhythm increase is related to heightened breathing, which is linked to fear-related 4 Hz oscillations in mice^34^. The remaining resonance peaks were exclusively of visual origin, as time-frequency analyses of responses to pulse and continuous luminance stimuli also showed increased power around resonance frequencies in theta (peak M_pulse_ = 6.97 Hz; M_ON-OFF_ = 7.72 Hz), beta (peak M_pulse_ = 15.19 Hz; M_ON-OFF_ = 15.94 Hz), and gamma bands (peak M_pulse_ = 36.1 Hz; M_ON-OFF_ = 39.84 Hz; Figure S5A,B). Taken together, similar to previous reports in humans, the temporal frequency tuning curves in mice displayed a banded structure with multiple resonance peaks (Figure 2D).

### Spatiotemporal responses beyond the retinotopic stimulus location

While responses to all flicker frequencies occurred at the retinotopic stimulus location of V1 and response amplitude decreased with distance from directly stimulated area, additional spatial peaks were present in response to some flicker frequencies (Figure 3A,C). Theoretically, such spatially non-uniform responses could result from structural anisotropies in anatomical connectivity across mouse visual cortex, or from propagation, interference and superposition of periodic stimulus-evoked traveling waves. Weak evidence for orientation anisotropy in the mouse visual cortex^35,36^ and the use of a uniform visual stimulus makes orientation anisotropy an unlikely explanation for spatial inhomogeneities in ssVER maps. To test the second hypothesis, we combined power and phase information in spatiotemporal pattern analysis and isolated the dominant spatiotemporal response features specific to each flicker frequency using generalized eigenvalue decomposition (GED; STAR Methods, Figure S4A). Although ssVER signal is typically captured by multiple GED components (Figure S4A,C)^37,38^, the first component explains the most variance (Figure S4C). Its associated spatial map, hereafter referred to as spatiotemporal response pattern, was used for further analysis.

**Figure 3.**
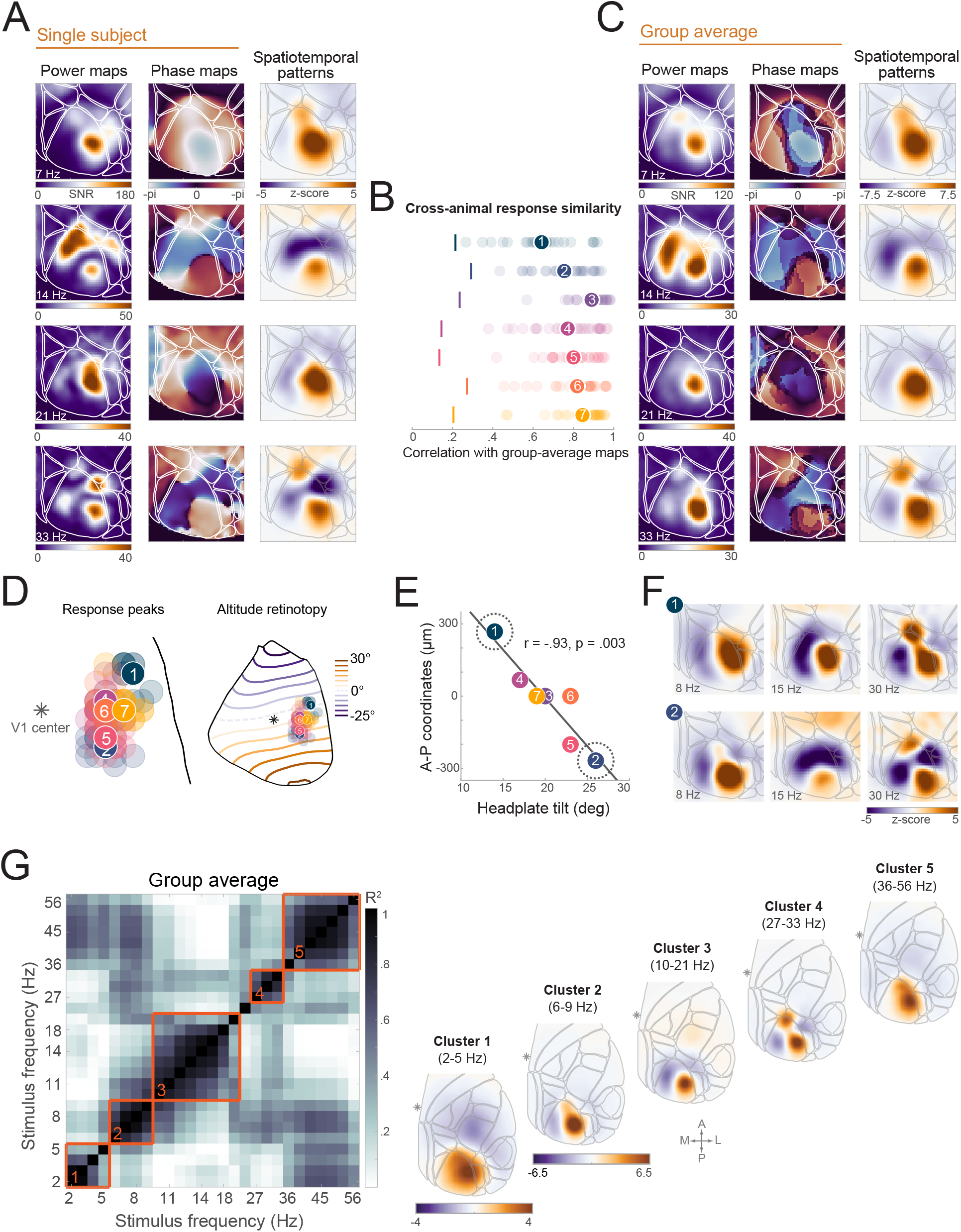
Spatiotemporal patterns in mouse primary visual cortex. (A) Exemplar flicker frequency-specific ssVER power maps, phase maps and spatiotemporal patterns (single-animal data). (B) Correlations between individual animal (panel A) and group-average (panel C) spatiotemporal response patterns in V1. Each row shows the data from an individual animal, with light-colored circles depicting correlation values for each 2-56 Hz flicker condition and dark-colored circles represent condition average. All correlations were significantly higher than those obtained in response to >56 Hz flicker conditions (vertical colored lines), which exceeded the temporal resolution of the glutamate sensor and where no correlation between animals is expected. (C) The same as A, but averaged across animals (N=7). (D) Left: Response peaks derived from condition-specific spatiotemporal patterns (light-color circles) and condition average (dark-color circles). Each animal is represented in a different color. Asterisk marks the center of V1 in Allen CCF. Right: Altitude contour plot of the mouse visual cortex from Zhuang et al. (2017) with overlaid average response peaks from each animal. Note that headplate position between animal 1 and 2 differed by ∼12^°^ and response peaks corresponded to ∼10^°^ of altitude difference in retinotopic map (contour lines are spaced at 5° intervals). (E) x-axis: headplate position relative to horizontal plane; y-axis: vertical response peak coordinates expressed relative to V1 center. (F) Exemplar spatiotemporal patterns from two animals showing extreme headplate positions (dashed circles in D). (G) Left: Group-average pairwise squared correlation matrix of spatiotemporal patterns. Using hierarchical clustering, five characteristic response patterns were identified. See also Figure S4.

Visual comparison of spatiotemporal response patterns with ssVER power maps revealed that secondary spatial peaks in power maps aligned with troughs in the spatiotemporal patterns (e.g. responses to 14 Hz and 33 Hz flicker in Figure 3A,C). Peaks and troughs in spatiotemporal patterns also coincided with antiphase regions in the phase maps, indicating opposite phases of stimulus-evoked wave at those locations in V1. Abrupt π-radian phase changes (phase reversals) separating large regions of a constant phase are characteristic of standing waves (Figure 3A,C). The region in which the standing waves formed was marked by near zero-amplitude contours in the power maps and roughly corresponded to the borders of the visual cortex. The correspondence between features in spatiotemporal patterns (first GED component) and power and phase maps, along with their variability across flicker frequencies, confirms that these patterns are not artifacts of the eigenvalue decomposition technique. While eigenvalue decomposition of spatially smoothed random noise returns spatial components ordered from low to high spatial frequency^39^, the spatial frequency of the first GED component in real data varied across flicker conditions (Figure 3A,C) and the second GED component had low spatial frequency (Figure S4D).

Despite previously reported substantial mouse-to-mouse variability in V1 size^40^, responses to different flicker frequencies were markedly consistent across animals (Figure 3A vs. C), with strong and significant correlations between single-animal and group-average spatiotemporal patterns in V1 (Figure 3B). As head position directly influences the viewing angle and retinotopic stimulus position, we investigated its impact on response variability across animals. We found a negative correlation between rightward headplate tilt and the spatial response peak in V1 along the anterior-posterior axis (r(5) = −0.93, p = 0.003; Figure 3E). A rightward headplate tilt led to a lower position of the left eye and a more posterior peak in V1 (Figure 3D), corresponding to a higher stimulus altitude based on retinotopic mapping^41^. Slight shifts in stimulus location rotated the overall spatiotemporal patterns, yet non-retinotopic secondary peaks persisted (Figure 3F), indicating that standing wave formation is invariant to stimulus location.

To investigate how ssVER cortical maps vary as a function of flicker frequency, we computed pairwise correlations between frequency-specific spatiotemporal patterns and performed hierarchical clustering on these correlation matrices. Clustering analysis was performed on both single-animal and group-average correlation matrices, with frequencies in the clusters constrained to be contiguous. At the group level, clustering revealed five characteristic ssVER cortical maps (Figure 3G). Per animal, on average 5.29 (SD = 0.76) clusters were identified, with the same five characteristic spatiotemporal maps present in all but one animal (four maps). In three subjects, a single group-level cluster was split into two sub-clusters (Figure S4B).

In summary, spatiotemporal pattern analysis indicated that spatial inhomogeneities in responses to some flicker frequencies resembled standing waves with a characteristic spatial scale and geometry that changed depending on the flicker frequency.

### Traveling and standing waves in mouse V1

Previous studies have demonstrated that visual stimuli elicit lateral spreading waves across V1^42– 46^, with some reporting wave deceleration at and reflection from V1 border^47^. To investigate wave propagation before standing wave patterns emerged, we analyzed responses to single pulses (Figure 4A-D,S5D). The spatiotemporal dynamics of single-pulse evoked responses closely resembled the first cycle 3–8 Hz ssVERs (slow enough flicker suitable for direct comparison with pulse stimulation; Figure 4A;S5C,D). A stimulus-evoked wave spread from retinotopic stimulus location within V1 and then propagated to medial and lateral visual cortices (Figure 4A; Movie 1,2). The first evoked response peak at the retinotopic stimulus location (M_peak_ = 50.5 ms; SD_peak_ = 5.2 ms) was followed by a widespread suppression across V1 (M_trough_ = 138.1 ms, SD_peak_ = 10 ms) and a second evoked response peak at the retinotopic stimulus location (M_peak_ = 219 ms; SD = 13.0 ms; Figure 4B,S5C). While evoked response peak time was identical across V1 (Figure 4B), waveform rise time increased with distance from the peak ROI, indicating an active wave propagation. In contrast, the falling phase occurred almost simultaneously across V1, suggesting spatially coherent suppression.

**Figure 4.**
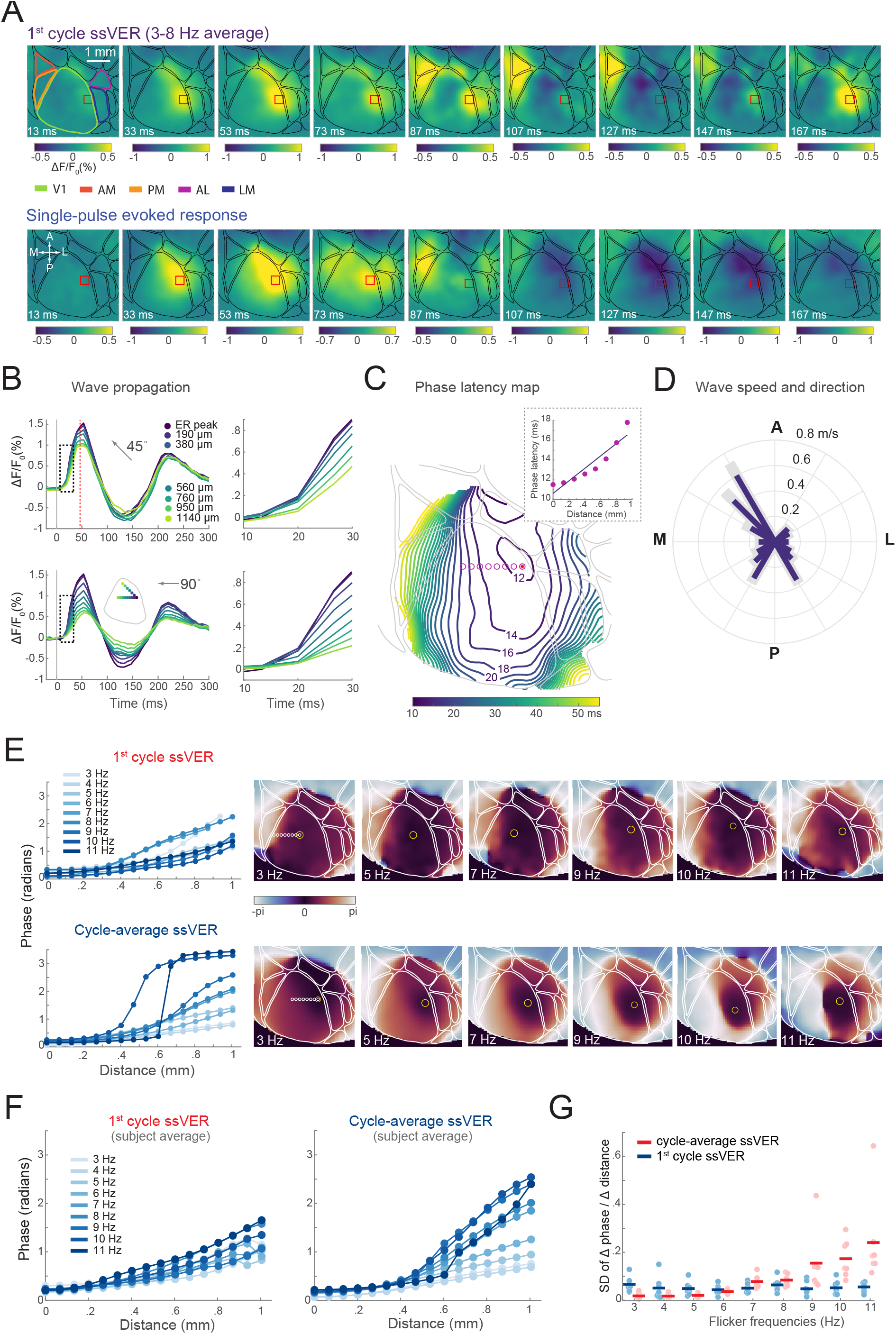
Traveling and standing waves in the mouse visual cortex. (A) Time-lapse images of stimulus-evoked activity during the first cycle flicker (3-8 Hz condition average; top row) and single-pulse stimulation (bottom row). The red rectangle marks the first evoked response peak location. Single-animal data. (B) Left: Evoked response waveforms from pixels along 45^°^ and 90^°^ directions (counterclockwise relative to the A-P axis) from the first response peak ROI. Right: Zoomed in plots showing delayed rise time with increasing distance from peak ROI. Higher amplitudes along the 45^°^ direction indicate active and faster wave propagation. (C) Phase latency map of evoked response to pulse stimulus, calculated +47 ms after the stimulus onset (reference timepoint), showing phase relation over space. The red dot marks the minimum (wave source) and isocontour lines depict phase offset (in ms) relative to the reference timepoint. (D) Group-average wave propagation velocity along different directions (blue bars). Grey areas indicate Mean + SEM. Velocity along some directions could not be estimated for all the animals (non-monotonically increasing phase at V1 borders) and therefore not plotted. (E) Left: Phase changes with distance along the 90^°^ direction starting from the minimum phase (yellow circle). Right: Phase latency maps of different flicker frequency ssVERs during the 1^st^ cycle (top row) and steady-state responses (bottom row). Single-animal data. (F) Same as E, but averaged across animals. (G) Standard deviation (SD) of phase change over a unit distance during the 1^st^ cycle (blue marks) and steady-state responses (red marks). Dots represent individual animals, horizontal lines mark group averages. A small SD indicates linear and high SD non-linear phase changes. See also Figure S5, Movies 1–4.

To characterize the stimulus-evoked traveling wave, we extracted the instantaneous phase from a band-pass filtered (5–25 Hz) signal at each pixel and constructed a phase latency map (phase latency differences relative to the timepoint preceding the first evoked response peak)^42^. Consistent with traveling wave behavior^44,48^, phase latency increased with distance starting from the wave source (minimum of the phase latency map; Figure 4C). Phase latency and cortical distance showed fairly linear relationship within 1000 μm radius (Figure 4C), with slight slowing near the visual cortex border, as indicated by the line density in the phase latency map (Figure 4C). We used the slope of a linear fit between phase latency and cortical distance relationship to compute wave propagation speed along different directions. The average speed propagation across all directions was 0.21 m/s (SD = 0.15 m/s; M_slow_ = 0.11 m/s; M_fast_ = 0.62 m/s), with some bias towards anteromedial directions (Figure 4D).

While a single flash evokes a single propagating wave, continuous rhythmic stimulation evokes multiple interacting waves that lead to steady-state spatial response patterns. Thus, we next analyzed the effects of flicker after a steady-state response has been reached (1000 ms post-flicker onset). To increase the signal-to-noise ratio, we computed peak-triggered average ssVER data, using every 5^th^ peak in the stimulus sine-wave. We compared phase changes over space during the first cycle of flicker stimulation with the steady-state response. In the 1st cycle, phase increased linearly from the retinotopic stimulus location for all flicker frequencies (Figure 4E,F), resembling a traveling wave evoked by pulse stimuli. In the steady-state response regime, phase maps varied with flicker frequency: As the flicker frequency increased, the maps progressively resembled a standing wave pattern characterized by abrupt phase changes (Figure 4E,F; Movie 3,4). These differences were captured by the main effect of condition (F(1,6) = 11.55, p = .015, 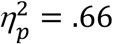), frequency (F(8,48) = 7.53, p < .001, 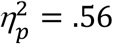), and significant two-way interaction (F(8,48) = 6.50, p < .001, 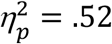) summarized in Figure 4G. Phase latency analysis was restricted to lower flicker frequencies due to temporal and spatial resolution of iGluSnFR in widefield imaging.

### Spatiotemporal responses: Linear wave equation solutions

Spatiotemporal patterns and phase-latency analysis indicated that cortical patterns in V1 induced by >5 Hz flicker exhibit standing wave characteristics (Figure 3,4). These standing waves formed within a roughly circular region outlined by the zero amplitude contours in ssVER power maps. Therefore, we investigated standing wave patterns that could theoretically form within a medium with isotropic connectivity and nearly circular boundaries.

The standing wave solutions to the linear wave equation are eigenmodes of the Laplace operator (∇^2^). We used a matrix form of discrete Laplace operator to represent the connections between pixels in the image. We assumed local (only adjacent pixels are connected) and isotropic connectivity within the modelled region and no activity at the boundary due to the observed small ssVER amplitude at the borders (Dirichlet boundary conditions). The modeled region was defined separately for each animal based on amplitude contours in ssVER maps (Figures 5B,S6). The geometric eigenmodes of the modelled region closely resembled the empirically observed spatiotemporal ssVER patterns evoked by different flicker frequencies (Figure 5A), supporting their interpretation as standing waves evoked by the rhythmic light stimulation.

**Figure 5.**
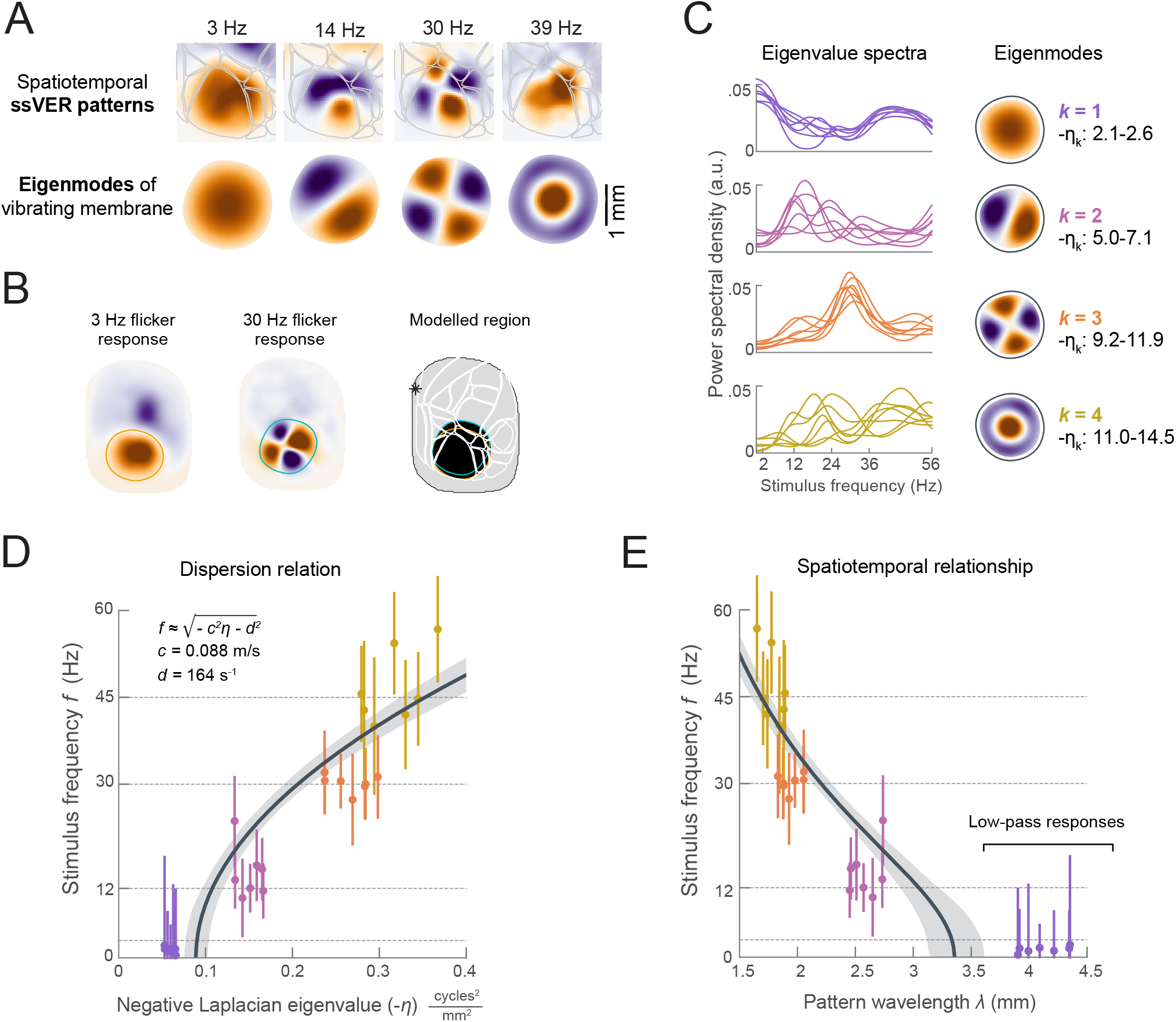
Relationship between flicker-induced cortical patterns and eigenmodes of vibrating membrane. (A) Visual similarity between flicker-evoked spatiotemporal patterns and standing waves (eigenmodes) of a nearly circular vibrating membrane. (B) The union of regions excited by 3 Hz and 30 Hz flicker was used to define a region on which standing-wave pattern formation was modeled. (C) Left: Eigenvalue spectra (y-axis) showing correspondence between each eigenmode and ssVER maps from different flicker conditions (x-axis). Eigenvalue spectra were computed as a squared dot product between eigenvectors corresponding to a specific Laplacian eigenmodes (*k = 1,2,3,4*) and normalized ssVER maps. Lines represent single-animal data. Right: Eigenmodes of the Laplacian operator with Dirichlet boundary conditions for a nearly circular modeled region, where *k* is the eigenmode index and is the range of Laplacian eigenvalues associated with each eigenmode. Eigenvalue depends on the geometry of the modelled region, which varied across animals. (D) Relationship between temporal stimulus frequency (*f*) and Laplacian eigenvalues corresponding to specific spatial patterns (eigenmodes). Color notation matches panel C and indicates different eigenmodes. Each data point represents a peak eigenvalue in eigenvalue spectra (panel C), with the vertical lines reflecting FWHM of the peak. The solid black line represents a regressed dispersion relation. (E) Relationship between the wavelength of the spatial pattern of each eigenmode and the temporal stimulus frequency *f*. See also Figure S6.

Next, we quantified the similarity between the geometric eigenmodes and the empirical spatiotemporal ssVER patterns (Figure 5C). Overall, we found that patterns evoked by the faster temporal frequencies were associated with higher spatial frequency eigenmodes, except for the fourth narrow “one-lobe” eigenmode which was not associated with a specific flicker frequency (Figure 5C). This mode overlapped with the area of retinotopic stimulation in V1 and captured a spatially localized response that was present at all flicker frequencies. Although eigenmodes were not uniquely associated with one specific flicker frequency, a range of flicker frequencies predominantly excited one specific eigenmode.

To determine the relationship between the temporal driving frequency and spatial frequency of ssVER patterns (dispersion relation), we assumed that the waves evoked by flicker travel at a constant speed *c* and are attenuated with distance *d* (damped wave equation). We hypothesized that the patterns evoked by flicker were consistent with standing-wave solutions to this wave equation, in which the stimulus temporal frequency *f* and the eigenmode *k* along with its associated eigenvalue *η*_*k*_ are square-root related 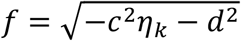 (STAR Methods). We used the eigenvalue spectra to identify a range of temporal stimulus frequencies *f* associated with particular Laplacian eigenvalue *η*_*k*_ and fitted the dispersion relation to obtain coefficients *c* and *d* (Figure 5D). The adjusted R^2^ of the fitted dispersion relation was 0.74. Wave propagation coefficient *c* on average was 0.087 m/s and corresponded to the lower end of empirically observed wave speeds (Figure 4D). The damping coefficient *d* was 164 s^-1^ (τ = 6 ms). Note that when *-c*^*2*^*η*_*k*_ *– d*^*2*^ *<* 0, the standing waves do not form and the spatiotemporal responses resemble a low-pass filtered version of the input. Assuming that the observed waves are planar, we derived relationship between the spatial scale of the evoked standing-wave patterns (wavelength *λ*) and flicker frequency (Figure 5E). Finer spatial response patterns were associated with faster flicker frequencies.

Overall, a damped classical wave equation effectively captured the phenomenology of large-scale neural responses to flicker. However, the complete picture likely includes richer nonlinear dynamics as, contrary to behavior of simulated neural networks (Rule et al., 2011), pattern-forming flicker frequencies (Figure 3G) were not identical to the temporal resonance frequencies (Figure 2C).

## DISCUSSION

Leveraging the high spatial resolution of widefield imaging with the fast kinetics of glutamate fluorescent reporter (iGluSnFR), we demonstrate that at a population level, mouse V1 follows much faster luminance changes than previously reported (56 Hz here, 24 Hz in Andermann et al.^49^). Stronger responses to some flicker frequencies were apparent as local resonance peaks in the temporal frequency tuning curves. Thus, mouse V1—just like human^5,8^ and cat V1^4,50^—favors specific temporal frequencies. We further found that responses to spatially localized flickering stimuli extended far beyond retinotopic stimulus location and were spatially inhomogeneous. By analytical means and modeling we demonstrate that the observed cortical spatial response patterns reflect standing waves resulting from propagation, interference and superposition of periodic stimulus-evoked traveling waves.

In mouse V1, response amplitude decreases as stimulus temporal frequency increases^1,13,51^, and slowly (1.5–3 Hz) drifting gratings elicit stronger responses on average^14,49,51^. While this suggests that V1 acts as a low-pass filter^1^, at a population level V1 can track flicker up to 40 Hz (in LFP recordings)^11,12^, with temporal frequency tuning curves from multiunit activity peaking at 6 Hz^13^. Using glutamate widefield imaging, we recorded significant responses for flicker frequencies up to 56 Hz and responses double the stimulus frequency were significant up to 66 Hz. Mouse ability to discriminate high frequency flicker has also been reported behaviorally, with critical flicker fusion frequency reaching 42 Hz^52^.

Confirming low-pass filtering characteristics of V1^1,13^, response amplitudes decreased with increasing flicker frequency. However, this decrease was not monotonic. After accounting for power-law scaling, we found resonance peaks in temporal frequency tuning curves in response to theta (∼5 Hz and ∼8 Hz), beta (∼15 Hz), and gamma (∼33 Hz) flicker. Similar to humans^8,9,53^, resonance frequencies in mice matched endogenous oscillations of the visual cortex in theta (∼4-8 Hz)^54^, beta (∼15 Hz)^54,55^, and gamma (∼33 Hz) bands^56,57^ (Figure S5A,B). The 3-5 Hz^33^ and 4-8 Hz^54^ oscillations likely originate from distinct sources. Yet, finer flicker frequency sampling (<1 Hz) is needed to confirm the separation of the ∼5 Hz and ∼8 Hz resonance peaks. Although resonance frequencies in mouse V1 were slower than those previously reported in humans^8^ (Figure 2D), the overall banded structure of temporal frequency tuning suggests that resonance phenomena are preserved across species and hallmark common cortical computations.

Consistent with previous reports^43,56,58^, both single pulse and flicker stimuli evoked traveling waves spreading from retinotopic stimulus location. The wave propagation velocity ranged from 0.1 to 0.6 m/s and was similar to those reported in awake ECoG recordings in mice (0.1-0.8 m/s)^56^, voltage-sensitive dye recordings in anesthetized cats (0.3 m/s)^44^, and awake primates (0.57 m/s)^42^. Such propagation speeds correspond to signal propagation along unmyelinated cortico-cortical horizontal fibers in superficial cortical layers^59^. While widefield imaging signal is dominated by activity in the superficial layers of the cortex^60^, glutamate release from thalamic nuclei projecting onto the superficial layers of V1 may also contribute to the measured signal^61^. Similar to previous reports^56,58^, stimulus-evoked waves propagated primarily in anteromedial direction. Projected onto visual space this direction corresponds to the direction of optical flow generated by the forward self-motion^62^, suggesting that stimulus-evoked waves follow a retinotopic path relevant for perception during locomotion. This direction may coincide with the direction of maximal information transfer^46^ and carry predictive information about the upcoming sensory input^63^.

Spatial response patterns to flicker frequencies >5 Hz extended beyond retinotopic stimulus location and exhibited standing wave characteristics within a nearly circular cortical region that included V1 and some higher visual areas (HVAs). Waves crossing V1 border were also evident in traveling wave analysis, consistent with the previous findings in mice^64^ and rats^47^. In physical systems, standing waves may form due to wave reflection from the boundaries. In the brain, boundaries can result from differences in connectivity patterns within and between cortical areas^41^. Based on connectivity patterns, mouse visual areas form two strongly interconnected modules, with comparably strong connectivity within V1 and between V1 and lateral visual areas^65^. This may explain why the standing waves were not confined only to V1 but also extended to HVAs. Although stimulus-evoked wave reflection occurs at architectonic V1/V2 border^47^, temporary functional boundaries could also contribute to standing wave formation. These temporary boundaries may result from interactions between stimulus-evoked excitatory wave and fast-spreading GABA_A_-mediated inhibition^45^ (Figure 4A; Movie 1), or between V1 feedforward waves and feedback waves from HVAs traveling at different speeds^56,66^. Regardless of their origin, cortical boundaries and geometry play a critical role in constraining wave-like dynamics across the entire brain^67^. We found that cortical standing waves induced by local flickering stimuli closely resembled the eigenmodes in a medium with isotropic connectivity and nearly circular boundaries. These standing waves provide insight into intrinsic functional architecture of the mouse visual cortex, highlighting the regions that are functionally connected and work synchronously.

The flicker-induced standing wave patterns observed here could explain frequency-dependent topographical variability of ssVEPs in human EEG^8,21^ and the apparent spatial organization in temporal frequency tuning reported in fMRI studies^18,19^. The double-peak topographies in EEG may reflect a standing wave. However, due to the spatial Nyquist frequency of EEG, only long wavelengths can be reliably measured and have been reported^20,21,68^. Thus, spatial response variability in macroscale signals like EEG could result from similar wave propagation and interaction principles reported here in the mouse visual cortex at a mesoscale.

In humans, flickering fields induce geometric hallucinatory percepts like moving circles, spirals, fans, lattices, and honeycombs^69^. The spatial frequency of these percepts increases with the flicker frequency^70^. According to neural field models, flicker-induced percepts are standing waves resulting from an interaction between external rhythmic input and endogenous network oscillations^22,71^. Rule and colleagues^22^ found standing wave patterns at stimulation frequencies that matched or were double the endogenous oscillations of the simulated network (resonance frequencies). Human empirical studies also show that percept formation speed and strength peak at resonance frequency of the human visual cortex (∼10 Hz)^72,73^. In this study, flicker frequencies evoking different spatial frequency standing wave patterns did not exactly match the temporal resonance frequencies (clusters in Figure 3G vs. peaks in Figure 2C). Although standing waves may contribute to entrainment and resonance, they are not necessarily linked to temporal resonance. Temporal resonance results from stimulation at frequencies matching endogenous oscillations of the corticothalamic network^4,13^, whereas standing waves arise from interference between periodically driven traveling waves supported by horizontal cortical connections^59^.

While suggestive, it remains unclear whether animals perceived standing wave activity beyond the stimulus area, and whether these patterns are related to flicker-induced geometric hallucinations in humans. The spatial frequency of standing waves in the mouse visual cortex was ∼1 mm for 30 Hz flicker, equivalent to ∼40^°^ of visual angle. In humans, geometric hallucinatory features are finer, spanning 1-2 mm of cortical surface^69^. Nonetheless, a positive correlation between standing wave spatial frequency and flicker frequency in mice aligns with previous findings in humans: Higher flicker frequencies induce finer and more detailed geometric hallucination and sudden doubling or halving of the flicker rate increases or decreases the number of hallucinatory features without altering the pattern^70,74^. While further experiments are needed, the standing wave spatial frequency could explain the temporal frequency-dependent spatial scaling of hallucinatory patterns.

In conclusion, using widefield imaging of a fast glutamate sensor in mice we empirically demonstrate the emergence of different spatial scale standing waves in the visual cortex in response to rhythmic sensory input and their relationship with temporal stimulus frequency.

## MATERIALS AND METHODS

### Animals

All experimental procedures were approved by Animal Welfare Committee of the University of Lethbridge and were performed in accordance with guidelines established by the Canadian Council for Animal Care.

Widefield glutamate imaging was performed in seven transgenic adult mice (5–6 months old, 4 females) expressing genetically encoded glutamate sensor iGluSnFR in glutamatergic neocortical neurons (Emx-CaMKII-Ai85 strain). The Emx-CaMKII-Ai85 mice were obtained by crossing the lines Ai85D (Jax026260)^75^, Camk2a-tTA (Jax007004)^76^, and Emx1-IRES-Cre (Jax005628)^77^. Prior to chronic window surgery, the animals were housed in groups of two to five under standard conditions in clear plastic cages with 12h light and 12h dark cycle. Mice had ad libitum access to water and standard laboratory mouse food. Following the implantation of a chronic imaging window, the animals were housed individually to prevent injuries and damage of the imaging window.

### Chronic Imaging Window Surgery

Chronic imaging windows over an intact skull were installed over the right hemisphere in 5–6 month old mice (4 female and 3 male), following previously published procedures^26,78^. The animals were anesthetized with 2.5% isoflurane, with anesthesia maintained with 1-1.5% isoflurane. Throughout the surgery, body temperature was monitored by a rectal thermometer and maintained at 37°C using a feedback-regulated heating pad. Following induction of the general anesthesia, the scalp was locally anesthetized with lidocaine (0.1 ml, 0.2%), after which the fur and the head skin was removed. Fascia and connective tissue were removed, exposing the right side of the skull. Thereafter, the clean and dry skull was covered with a clear version of C&B-Metabond dental cement (Parkell, Inc.) and a metal head-plate was glued using dental cement. Finally, the skull was covered with a glass coverslip. Following awakening from anesthesia, the animals were put in the recovery room for a 24h period, after which they were placed in their home cage and monitored daily. The animals were allowed to recover from the surgery for at least 1 week before habituation to the experimental procedures was started.

Analysis of pilot recordings revealed that despite extensive measures aimed at protecting the imaging area from exposure to the white LED visual stimulation device (custom-designed 3D printed plastic protection and black non-reflective curtain covering the imaging area, see Figure 1), light leaks through the skin and small openings between the head-plate and the skull were present in the frontal part of the imaging window (Figure S1E). Therefore, 3 weeks after the initial surgery, the animals underwent a short procedure during which an additional layer of black dental cement was applied over the frontal and lateral parts of the head-plate fixation and the skin. This procedure was performed under general anesthesia, controlling the body temperature and using the antiseptic techniques as previously described. This additional procedure successfully eliminated light artifacts (Figure S1F).

### Habituation

Following a full recovery from the chronic imaging window implantation surgery, the animals were trained to remain head fixed using a 7-day incremental training time procedure, which minimizes animal stress. The first day, animals were placed on the head fixation setup for 5–10 minutes and were allowed to freely move and explore the setup. On subsequent days, the mice were head-fixed using two clamps for 5 minutes, with increasing time per session (from 5 minutes up to 30 minutes), food reward followed each training session. During habituation and recording, body movement and stress were minimized by putting them in the enrichment plastic tube from their home cage. Sounds and vibrations in the recording setup and the room were minimized, light stimulation was present during the habituation stage to simulate the experimental protocol.

### Imaging

In vivo glutamate imaging was performed in awake head-fixed animals using a charge-coupled device (CCD) camera (1M60 Pantera, Dalsa, Waterloo, ON) and an EPIX E8 frame grabber with XCAP 3.8 imaging software (EPIX, Inc., Buffalo Grove, IL). The onset and offset of CCD camera and behavioral camera, onset and offset of each frame, as well as voltage changes corresponding to luminance changes of the visual stimulation device were recorded in Clampex (Molecular Devices, Sunnyvale, CA) and used for data analyses to determine frames during which visual stimulation was delivered. During the imaging, the behavioral camera and an infrared light were placed in front of the animal to constantly monitor active behavior and any signs of distress.

To capture the fast glutamate sensor iGluSnFR kinetics, images of iGluSnFR activity were captured at 150 Hz sampling rate. The glutamate fluorescent indicator was excited using blue LED light (Luxeon K2, 470 nm), which was band-pass filtered using an optical filter (Chroma

Technology Corp, 467-499 nm). Light emitted from excited fluorescent indicators was passed through a band-pass optical filter (Chroma, 510 to 550 nm; Semrock, New York, NY) and a macroscope composed of front-to-front optical lenses. The focal length of the lenses was adjusted such that the field of view was 8.6 × 8.6 mm (128 × 128 pixels, with 67 μm per pixel). To minimize the effect of hemodynamic signal originating from large cortical blood vessels, we focused the optical lens at ∼1 mm depth.

### Visual stimulation setup

We used a custom-built setup of white light-emitting diodes (LEDs; luminous intensity 6900 mcd, color temperature 9000K, Model C513A-WSN, Cree Inc.) to deliver visual stimulation (the setup previously used in Gulbinaite et al.^8^). The LED array was placed in a cardboard box, with a circular aperture subtending 35° of visual angle covered by a sheet of tracing paper (6 cm away from LED array). The visual stimulation box was placed on the left side of the animal, 8 cm from the left eye, at a 50° angle to the animal’s body axis. The luminance of white LEDs was linearized using a quadratic function, such that a linear increase in voltage corresponded to a linear increase in luminance (measured in cd/m^2^). Flicker was implemented as a sine-wave modulation of the power supply to the LEDs (luminance changes from 0 to 215 cd/m^2^), which was controlled using a microcontroller of the STM32F4-Discovery board by adjusting the width of pulse-width modulated (PWM) signal and linearizing the current using a low-pass filter. MATLAB was used for communication with the STM32F4-Discovery board via a USB virtual com port protocol (programmed in C++). Voltage changes driving the changes in luminance of stimulation LEDs were recorded in pClamp (Molecular Devices, Sunnyvale, CA) as a separate channel in Axon Binary Files (ABFs). Recorded waveform was used to determine the CCD camera frames during which visual stimulation was delivered.

### Stimulation protocols

#### Steady-state visual evoked potentials

On each trial, the animals were exposed to 10 s of flicker followed by 10 s inter-trial interval of complete darkness (Figure 1B). Mice were exposed to flicker frequencies that ranged from 2 to 72 Hz (logarithmically spaced 30 different frequencies). For all frequencies, the stimulation started and ended with maximal luminance (215 cd/m^2^, π/2 phase of the sine-wave cycle). Each imaging session lasted up to 30 minutes, during which each flicker frequency was presented twice in pseudo-random order. Each animal underwent 5 experimental sessions (i.e. 5 recordings on separate days). Thus in total, each flicker frequency was presented 10 times (10 s × 10 = 100 s per frequency), and the recording duration was comparable to that previously used in a human study (96 s per flicker frequency)^8^.

After each experimental session, the excitatory blue LED light used to excite the glutamate fluorescent indicator was switched off and 5 Hz flicker was presented for 3 trials (10 s flicker, 10 s inter-trial interval). This control recording was used to determine the presence and the extent of any potential light artifact from the visual stimulation leaking into the imaging window through the skin and/or junction of the 3D printed light-blocking cone and the head-plate (see Figure 1A,S1E).

#### Visual evoked-potentials

In addition to sine-wave stimulation, we recorded responses to single light pulses (20 ms) and continuous light stimulation (5 s) using 1500 cd/m^2^ luminance stimuli to characterize spatiotemporal dynamics in response to single stimuli and on/off responses.

#### Sensory evoked-potentials

Stimulation of other senses was performed in anesthetized animals (1% isoflurane) and was used to determine the coordinates of other primary sensory areas^26^: Hindlimb area of primary somatosensory cortex (SSp-ll), forelimb area of the primary somatosensory cortex (SSp-ul), primary barrel sensory cortex (SSp-bfd), and primary auditory cortex (AUDp). The forelimb and hindlimb were stimulated electrically (1 mA, 1 ms electrical pulse) with a thin acupuncture needle inserted into the paws. The C and D whiskers were stimulated using a piezoelectric bending actuator, and a single 1 ms tone (25 kHz) was used for auditory stimulation. Each stimulus was followed by 10 s inter-stimulus interval. Sensory evoked potentials were calculated by averaging 20–30 trials.

### Image preprocessing and analysis

#### Spatial co-registration across recordings and to the Allen Mouse CCF

Recordings obtained on different days and different animals were co-registered using rigid transformations (translation and rotation). Prior to each recording session, a stack of 15 reference images was taken by focusing the lens around the depth of blood vessels. Reference images were then merged into one image using ImageJ. The transformation matrix used for the alignment across recording sessions was computed as follows. First, the images from each animal’s first session were translated (x and y direction) to match bregma (marked during the window-placement surgery) to an arbitrarily chosen reference point (x=10, y=30), and rotated such that the axis connecting bregma-lambda was parallel to the bregma-lambda axis in the Allen Institute Common Coordinate Framework (CCF) map. Thereafter, reference images from each subsequent session were co-registered to the reference images of the first acquisition using surface vasculature and anatomical landmarks. Hereby obtained transformation matrices were then applied to the image stack of a corresponding recording session. 2D Allen CCF map was obtained from 3D Allen CCF mouse brain rotated 30^°^ counterclockwise around the anterior-posterior axis to accommodate for the tilted (∼30^°^) placement of the headplate over the right hemisphere. The quality of alignment to Allen CCF was verified by creating a functional map: Plotting the evoked potentials from all sensory stimulation paradigms (Figure 1C,S1B).

#### Preprocessing and hemodynamic artifact removal

Continuous recordings were first epoched around the stimulus onset (−2–15 s for ssVER paradigm, -2–9 s for 5 s continuous visual stimulation, -2–4 s for visual pulse stimulation, -1–1 s for auditory, forelimb, hindlimb and whisker stimulation). Non-brain pixels were masked prior to further signal preprocessing steps. To remove temporal changes in glutamate fluorescence related to hemodynamic response, breathing and heart rate that are inherent to imaging green fluorescent protein indicators^24^, we applied global signal regression (GSR). If we denote the fluorescence signal at each pixel *i* and timepoint *t* as *S*_*i*_*(t)*, the global signal regressed out is a time-average of each pixel:

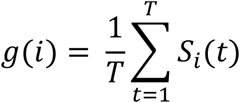

The GSR-corrected signal is then:

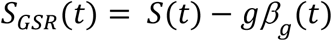

where *β*_*g*_*(t)* = *g*^*+*^*S(t)* and *g*^*+*^ is a pseudo-inverse of *g* (*g*^*+*^ = [*g*^*T*^*g*]^-1^*g*^*T*^). Such signal normalization over time effectively removes hemodynamic artifacts in pixels containing blood vessels and preserves the real brain signal in response to visual stimulation used here. Changes in glutamate fluorescence of each pixel at each time point were expressed as percent change relative to pre-stimulus baseline window -1000 to -100 ms (ΔF/F_0_ × 100%, where F_0_ is the average fluorescence signal during the baseline period, and F is fluorescence at each time point, ΔF = F-F_0_).

Application of GSR allowed to remove 8–12 Hz heart-rate related hemodynamic signals which result from the overlapping excitation and emission spectra of green fluorescent proteins and absorption spectrum of hemoglobin^28^ (Figure 1D-E). However, GSR only partially attenuated a slow, BOLD-like wave that was locked to the stimulus onset. Slow waves starting around 1 s post-stimulus and lasting for 4-5 seconds have been reported before and are a result of neurovascular coupling^1,28^.

To correct for the slow neurovascular response superimposed on evoked responses to single pulse and ssVERs (sine-wave stimulation), we performed *robust detrending*: Iteratively fitting the *n*^th^ order polynomial to the data and subtracting it^29,30^. Robust detrending effectively removed all slow trends unrelated to experimentally controlled visual inputs. Given that detrending is highly sensitive to sharp amplitude changes in the timeseries such as ERPs (here, a response to flicker onset), fitting of which can introduce spurious fluctuations in the detrended signal or even partially removal of a neural signal^29,30^, we performed polynomial fitting excluding the first 100 samples (667 ms) after the stimulation onset and offset (for similar approach see van Driel et al.^30^). To perform robust detrending we used *nt_detrend* function from NoiseTools MATLAB toolbox (http://audition.ens.fr/adc/NoiseTools/)^29^. Robust detrending is an iterative procedure, such that a time window in each pixel timeseries is fitted with a high-order polynomial, and residuals between the fit and the raw signal are computed. The fit is then subtracted from the raw signal and time points at which residuals surpass a predefined threshold (3 standard deviations from the mean) are downweighed (weights set to 0) and are not fitted in the next iteration. The maximum number of iterations was set to 3 (default value).

By setting the initial weight vector to 0 during the 0–700 ms window relative to the stimulus onset and offset (in case of flicker and continuous stimuli), we made sure that the real neural signal was not removed by robust detrending procedure. This window was chosen based on: (1) the duration of an evoked response in mouse V1 reported using voltage sensitive dye imaging^78^, which is not affected by hemodynamics^1^ and (2) the onset of to-be-fitted slow wave (negative ΔF/F_0_ values started around 500 ms after flicker onset). The higher the order of the polynomial, the smaller the fluctuations that are fitted and subsequently subtracted. Thus, there is a tradeoff between artifact and real signal removal. We empirically tested polynomials with order 10, 20, 30 and 50, and selected the order 20 polynomial, because evoked response peak amplitudes were not altered after polynomial removal and computation time was acceptable for these large datasets. The polynomial fitting was done in sliding windows of 100 frames (667 ms), with 50% window overlap.

#### ssVER power and phase calculation

ssVERs are assumed to be stationary narrow-band signals and ssVER power spectra are typically calculated from trial-average waveforms. However, fluctuations in instantaneous ssVER frequency indicate cycle-to-cycle variations in response waveforms which would be lost after averaging^8^. For this reason, FFT was performed on single-trial data in the 1000–10000 ms time window (relative to the stimulus onset) and zeropadded to obtain 0.1 Hz resolution for power spectra. The absolute value of FFT coefficients were squared and averaged across trials to obtain ssVER power spectra. When calculating power spectra, the first 1000 ms post-stimulus were excluded, as it contains evoked response related to the flicker onset and steady-state response takes time to develop^8^. To facilitate comparison of ssVERs generated in response to different flicker frequencies and to account for 1/f power-law scaling effect (stronger endogenous brain activity at slow frequencies)^79^, power spectra from each flicker condition were converted to SNR units^16^. This is achieved by expressing each datapoint in the raw power spectrum as a ratio between the power at the frequency of interest and the average power at the neighboring frequencies (±1 Hz) of the FFT spectrum, excluding 0.5 Hz around the frequency of interest (see Figure 1SC for the illustration of the procedure):

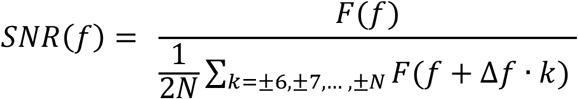

where *Δf* = 0.1 Hz, *N* = ± 1 Hz (10 bins). We constructed temporal frequency tuning curves by plotting response power at the flicker frequency from the power spectrum of response to each flicker condition. The frequency resolution in temporal frequency tuning curves depended on the number of flicker frequencies. We identified peaks in the temporal frequency tuning curves using MATLAB *findpeaks* function. Bimodality in the theta band of temporal frequency tuning curve was assessed using Hartigan’s dip test^80^, where power from 3–14 Hz flicker conditions from each animal was used to build an ordered distribution. The empirically observed dip value was compared to 1000 bootstrapped dip values to determine significance.

We also used FFT analysis on trial-average data to obtain phase maps by computing the angle of complex Fourier coefficients at each flicker frequency of interest for each pixel. In this case, we used trial-average data because of higher signal-to-noise ratio for phase estimation.

#### ssVER spatiotemporal pattern analysis

Spatially variable patterns in average power and phase maps indicated that the spatial distribution of ssVERs depended on flicker frequency. However, power and phase maps derived from FFT over the entire trial window discard temporal amplitude fluctuations and provide only a static view. Thus, in the follow up spatiotemporal pattern analysis we combined temporal and spatial properties of ssVERs by narrow-band pass filtering the data around the flicker frequency and using two-stage generalized eigenvalue decomposition (GED) on covariance matrices (detailed illustration of analysis steps is depicted in Figure S4A). The end goal of this procedure was to compute a set of pixel weights (empirical spatial filters) that maximally separate the signal data (response to specific flicker frequency) from the reference data (brain response to spectrally adjacent non-flicker frequencies). Given the high number of dimensions (∼8000 pixels containing brain data), GED can become numerically unstable^38^. Therefore, we first compressed the data by projecting out principal components that explained less than 0.01% of total variance. The resulting compressed data were reconstructed as follows:

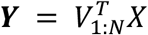

where **V** are principal eigenvectors, **X** is the data matrix with shape pixels × time × trials, and N is a number of retained components.

In the second step, we performed generalized eigenvalue decomposition (GED) to find spatiotemporal patterns in pixel covariance matrices that are characteristic to specific temporal frequency response. This was implemented by finding a linear weighted sum over all brain pixels that would result in the highest multivariate signal-to-noise ratio. The “signal” (**S**) here is the response to flicker frequency, and “noise” or “reference” (**R**) is the signal at spectrally adjacent non-flicker frequencies. In practice, this was done by eigenvalue decomposition of the matrix product R^-1^ S and solving the equation:

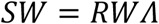

where **S** and **R** are covariance matrices, **W** is the matrix of eigenvectors and **Λ** is a diagonal matrix of eigenvalues.

The **S** and **R** covariance matrices were constructed as follows. First, data time series **Y** were narrow band-pass filtered using Gaussian filters: (1) centered at the flicker frequency *f* with FWHM = 0.5 Hz; (2) filter centered at *f -* 2 Hz with FWHM = 2 Hz; and (3) filter centered at *f* + 2 Hz with FWHM = 2 Hz. Data filtered at the flicker frequency is further referred to as “signal” (S), and data filtered at the neighboring frequencies is referred to as “reference” (R). Single-trial temporally-filtered data from 1000–10000 time window (relative to flicker onset) were concatenated and were used to compute *N* × *N* covariance matrices: one **S** matrix and two **R** matrices that were averaged. When computing covariance matrices, the first 1000 ms after flicker onset were excluded, because the quality of GED spatial filters (columns of the matrix **W**) is poorer when sharp transient responses to flicker onset are included^37^. Covariance matrix **R** was shrinkage regularized by adding 1% of its average eigenvalues (0.01·mean[eig(**R**)]) to the diagonal of the **R** matrix prior to GED^38^. Shrinkage regularization reduces the influence of noise on the GED, and improves stability and generalization. GED performed on covariance matrices **S** and **R** returned a matrix of eigenvectors (**W**) and a diagonal matrix of eigenvalues (**Λ**). Eigenvector **w** (spatial filter) associated with the highest eigenvalue λ was multiplied by “signal” covariance matrix **S** to extract the spatial response pattern characteristic to a specific flicker frequency, and reconstructed back to pixel space: 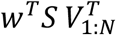. The first component also explained the most variance, which was computed as a ratio between eigenvalue associated with each component and the sum of all component eigenvalues (Figure S4C). Note that GED and principal components analysis (PCA), though both built on eigendecomposition, have different assumptions and constraints, and give different results^38^. We used GED to isolate and denoise existing patterns in the data, which is an application for which PCA generally performs poorly^37^.

#### Wave propagation analysis

To quantify and visualize stimulus-evoked traveling and standing waves we used the phase latency method (https://github.com/mullerlab/wave-matlab)^42^. To increase the signal-to-noise ratio, we first re-epoched ssVER data relative to every 5^th^ peak in the stimulus sine-wave (excluding the first 1000 ms containing flicker-onset evoked responses) and averaged these short epochs. Thereafter, we narrow band-pass filtered the data using Gaussian filters centered at the flicker frequency with FWHM = 1 Hz. Evoked responses to single-pulse stimulation were band-passed filtered (5-25 Hz) using forward-reverse fifth-order Butterworth filter. After filtering the ssVERs and single-pulse evoked responses, we extracted the instantaneous phase at each pixel using a Hilbert transform. Thereafter, we computed the phase offset (in milliseconds) relative to the timeframe prior to the first evoked response peak^42^. This ensures that phase latency maps represent phase relationships over space at the timeframe of ssVER or evoked response peak. The minimum of the phase latency map was taken as the wave source. For ssVER data, phase latency maps were computed only for low-frequency conditions (<13 Hz), as high-frequency flicker stimulation rendered phase-based analyses impossible. Phase latency maps were smoothed only for representation purposes (robust spline smoothing^81^), unsmoothed phase latency was used for wave velocity calculations.

To compute the wave velocity, we performed a linear fit to the phase latency and distance (within 1000 μm radius from the wave source) relationship along 16 directions (0, 30, 45, 60, 90, 120, 135, 150, 180, 210, 225, 240, 270, 300, 315, 330 deg). The inverse of the linear fit slope was taken as a wave propagation speed. For each animal, we only considered directions along which phase latency was monotonically increasing from the wave source. For some animals, wave speed for directions around anterior-posterior axis (90, 120, and 270 deg) could not be computed due to close proximity of the wave source to the V1 border and thus these directions are not represented in the group average plot in Figure 4D.

### Statistical analyses

#### Evaluating statistical significance of ssVERs

To evaluate statistical significance of ssVER responses, for each flicker condition the power spectra were calculated using trials on which neither the flicker frequency nor harmonically related frequencies were presented. This allows to estimate expected response SNR (“noise” SNR distribution). On average, “noise” SNR distribution was built from 261 trials (SD = 25.37). The 95^th^ percentile of “noise” SNR distribution is plotted in Figure 2C and referred to as “H0 hypothesis”.

#### Cortical map statistics

To evaluate and compare the spatial extent of responses across different flicker frequencies, power maps at the flicker frequency (“signal”) were compared against adjacent conditions when that flicker frequency was not present (“noise”). For example, 7 Hz ssVERs signal (10 trials) was compared against 7 Hz activity elicited when stimulus flickered at 6 Hz and 8 Hz (20 trials in total). Three different techniques were used to generate null-hypothesis (H0) distribution: (1) drawing “signal” and “noise” condition trials by resampling with replacement (bootstrap) from all trials pooled together^82^; (2) resampling without replacement (permutation testing) by permuting all the trials and randomly assigning a condition label^83^, and (3) bootstrap-t technique, in which the two conditions are mean-centered and bootstrapped samples are drawn from the two conditions separately. Given the statistical robustness of bootstrap-t approach for small sample sizes, independence of each draw and almost identical results of all three resampling approaches, results on bootstrap-t are reported here. The procedure was performed as follows.

First, cortical maps of “signal” trials and “noise” trials were z-scored, and two-sample t-test performed at each pixel to obtain the *observed* t-values. Thereafter, signal and noise trials were mean-centered. Second, two groups of trials were separately sampled with replacement and a t-test performed for each bootstrap (1000 iterations), thus building a distribution of *bootstrapped* t-values. Third, the observed t-values were z-scored using the mean and SD of bootstrapped t-values. Fourth, SNR differences between conditions were considered statistically significant if the z-scored t-value at each pixel in the cortical map was larger than 3.0902 (*p* < 0.001). Finally, cluster-based correction was applied to correct for multiple comparisons over pixels, such that only contiguous pixel islands (clusters) that were larger in size than 99% of H0 cluster size distribution were considered significant. The H0 distribution of cluster sizes was obtained by thresholding cortical maps from each of the 1000 iterations at p = 0.01, and storing the maximum cluster extent (sum of t-values within a cluster). Clusters of contiguous pixels in the observed t-value cortical maps were considered significant if the cluster size was bigger than expected under the null hypothesis. To obtain more stable estimates of cluster significance, we ran a “meta-permutation test” by repeating pixel-level and cluster-level permutation procedure 20 times. Thus, the average of 20 real z-scored t-values and the average of 20 cluster thresholds were used.

#### ssVER spatiotemporal patterns

To identify similarities across cortical maps elicited by different flicker frequencies, we computed pairwise correlations between frequency-specific spatiotemporal patterns identified using a two-stage GED (see subsection *ssVER spatiotemporal pattern analysis*). These correlation matrices, a prominent feature of which is a block diagonal form, were submitted to hierarchical clustering which allowed to group ssVER cortical maps and identify characteristic spatiotemporal patterns. To facilitate clustering, the correlation matrices were squared. To build a hierarchical tree (dendrogram), we performed agglomerative clustering using Ward’s minimum variance criterion which minimizes the total within-cluster variance (Matlab function *linkage*). The hierarchical tree was then pruned by setting the number of lower branches (leaf nodes) to maximum 4, which was determined empirically and allowed to distinguish 4-6 clusters of spatiotemporal patterns per animal. Frequencies in the clusters were constrained to be contiguous, as we were interested to examine changes in spatiotemporal response patterns with increasing flicker frequency. Note that clustering is an ill-posed problem in machine-learning, and different clustering algorithms and parameters may lead to somewhat different results. We applied our algorithm consistently to all data to avoid introducing systematic biases, but it is possible that the exact cluster boundaries would change with different analysis methods.

### Modeling standing waves

#### A linear wave equation

As noted in the main text, we explored whether empirically observed standing wave patterns could be approximated by d’Alembert’s classical wave equation with a damping term:

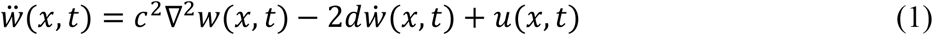

where *w*(*x, t*) are the observed spatiotemporal wave responses at time *t* and cortical position ***x*** = {*x*_1_, *x*_2_}, and *∇*^2^ is the spatial Laplacian operator which sums the second spatial derivatives in *x*_1_ and *x*_2_. The single dot above *w* denotes the first derivative with respect to *t*, and the double dot the second derivative. The discrete Laplacian in 1D corresponds to convolution with the kernel {1, −2, 1}, where a “restoring force” is proportional to the wave’s displacement at each location *x* relative to its neighbors. We included a viscous damping force 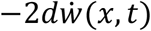 that attenuates the wave response over time. Parameter *c* is related to the wave propagation velocity and *d* to the wave damping. We chose 2*d* rather than *d* to simplify the expression for dispersion relation (see the next section “*Dispersion relation*”). We modeled the flickering stimulus as a spatiotemporally separable product of a spatial component *u*(*x*) with temporal modulation *u(t)*, i.e. *u(x, t)* = *u*(*x*) *u(t)*. Parameters *c* and *d* were determined using a fitting procedure (see section “*Estimating dispersion relation*”).

#### Dispersion relation

A linear wave equation such as in equation (1) makes specific predictions about the relationship between the temporal frequency of the flickering stimulus and evoked spatial modes (a dispersion relation). The spatial component of separable solutions to a linear wave equation are eigenmodes of the Laplace operator ∇^2^. Each eigenmode has a characteristic spatial and temporal frequency, and indicates which standing wave patterns can be expected to form at various temporal stimulation frequencies.

To derive a dispersion relation and compute the eigenmodes, we performed a change of basis to diagonalize equation (1). We used the two-sided Laplace transform for the temporal basis, expressed as: *ξ*(*s*) = *∫ w*(*t*) *e*^-*st*^ *dt*, which is related to the Fourier transform in angular frequency *ω* by the substitution *s ← iω*. The spatial basis consists of eigenfunctions *ϕ*_*k*_(*x*) (indexed by *k*) of the Laplacian operator ∇^2^. These eigenfunctions satisfy the eigenvalue equation *η*_*k*_*ϕ*_*k*_(*x*)= *∇*^2^*ϕ*_*k*_(*x*), with eigenvalue *η*_*k*_. On an infinite (or periodic rectangular) domain, these are sinusoidal plane waves. Here, however, the domain was a nearly circular patch of cortex, and Laplacian eigenmodes with Dirichlet boundary conditions were solved numerically (see section “*Defining region of cortex for modeling spatial eigenmodes*”).

Switching to the Laplace domain in time (*s*) and the Laplacian eigenfunction spatial basis (*k*), equation (1) becomes:

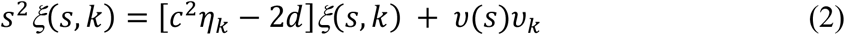

where *v*(*s*) and *v*_*k*_ are the projections of the input’s temporal and spatial components onto Fourier-Laplace temporal basis and spatial Laplacian eigenspace, respectively. Collecting terms in (2) and solving for *ξ(s,k)* gives a transfer function *g(s,k)* that maps stimulus input (***υ***) to the spatiotemporal wave response in cortex (***ξ***):

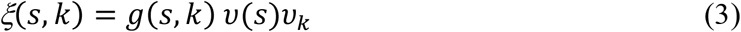

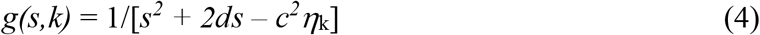

The system’s modes are given by the poles of this transfer function, i.e. when the denominator of *g(s,k)* is zero. Solving for *s*, we get:

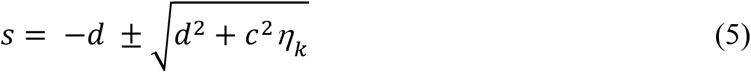

Oscillating solutions appear when *s* = −*d* ± *iω* is complex, which occurs when *d*^2^ *+ c*^2^*η*_*k*_ *<* 0. The real part of these solutions, −*d*, determines how quickly modes decay in time. The imaginary component determines the oscillation frequency *ω* (radians/second), and solutions always appear in conjugate pairs with a positive and negative *ω*:

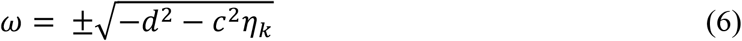

By squaring both sides of the equation (6), we obtain a relationship between Laplacian eigenvalue *η* and the square of the temporal frequency *ω*:

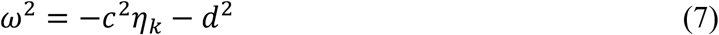

#### Spatial-scale interpretation of Laplacian eigenvalues

In Figure 5E, we present an intuitive interpretation of the Laplacian eigenvalues (*η*_*k*_) in terms of the spatial wavelength of plane-wave solutions to the wave equation with no boundary. On an infinite, unbounded domain the Laplacian eigenfunctions *ϕ*_*k*_*(x)* are Fourier modes with frequency components for each spatial direction *v* = {*v*_1_, *v*_2_}, and eigenvalues *η*_***v***_ = −∥***v***∥^2^, where ∥***v***∥ is the net spatial frequency of the wave. By analogy, eigenfunction solutions on the bounded domain can be written in units of spatial angular frequency (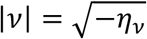 in units of radians/mm) or wavelength (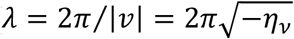 in units of mm per cycle). By solving wavelength λ in *η*_*v*_ = −(2*π*/*λ*)^2^, we obtain relationship between the wavelength *λ* of the spatial pattern and flicker frequency *ω* (Figure 5E). We report *λ* in mm units and *ω* in Hz units.

#### Defining region of cortex for modelling spatial eigenmodes

Based on visual inspection, spatiotemporal patterns extracted using two-stage eigenvalue decomposition elicited by 3 Hz and 30 Hz flicker stimuli contained spatially well-defined near-zero amplitude contours. We used the union of regions excited by these two flicker conditions to define the excited region of posterior cortex. The region was defined separately for each animal. The extracted regions were roughly circular, spanning 3.9–4.3 mm along the anterior-posterior axis, and 3.7–4.4 mm along the mediolateral axis (Figure S6). Alignment with the Allen CCF indicated that this region includes the extended visual cortex (V1 and several higher-level visual areas).

#### Calculating Laplacian eigenmodes

Calculation of Laplacian eigenmodes requires to define how traveling waves interact with areal boundaries. Given the phase reversal in phase maps and near-zero ssVER amplitude near the boundary, we assumed Dirichlet type boundary conditions of the modelled region. We constructed a matrix representing a discrete Laplace operator on the region of posterior cortex excited by the flickering stimuli (a region of 55–65 pixels width and height, containing 3306–4160 pixels each). For this, we first constructed an adjacency matrix (**A**), with rows and columns indexed by pixel location, with the (*i, j*)^th^ element set to 1 if pixel *i* and *j* are adjacent, and 0 otherwise. The discrete Laplace operator matrix is then defined as **L = A**−**D**, where **D** is a diagonal matrix. We divided the resulting matrix **L** by the squared grid spacing (223 mm^−2^), giving this matrix physical units of a second spatial derivative. To implement Dirichlet boundary conditions, we dropped rows and columns from **L** corresponding to pixels outside the specified boundary. We used sparse matrix formats and scipy’s sparse linear algebra routines to calculate the eigenvectors *v*_*k*_ and eigenvalues of *η*_*k*_ of this discrete Laplacian matrix (scipy.sparse.linalg.eigsh). Exemplar eigenmodes from one animal are shown in Figure 5A and are compared to spatiotemporal patterns extracted using two-stage GED procedure.

#### Eigenvalue spectra

We quantified the extent to which each Laplacian eigenmode was triggered by a particular flicker frequency *ω* by computing the similarity between the two cortical maps. At each pixel ***x*** we performed FFT on single-trial data and extracted the complex Fourier coefficients ***z***_*ω*,***x***_ at the stimulus frequency *ω*. We normalized each ***z***_*ω*,***x***_ to unit length 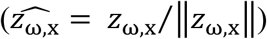 to account for the power-law scaling (decrease in response amplitude with increased frequency). We then calculated the variance in 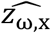 explained by eigenmode *v*_*k*,***x***_ by computing the squared dot product 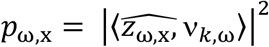. This way we obtained eigenvalue spectra: One for each eigenmode (Figure 5C). The eigenvalue spectra were smoothed with a Gaussian kernel (σ = 5 Hz). In Figure 5C, we report normalized eigenvalue spectra for each eigenmode in units of power density per Hz.

The approximately circular 2D domain contains eigenmodes with similar eigenvalues. For example, the “four-lobe” pattern appears as two eigenmodes, with one rotated by 45°relative to another. Focal stimulation will excite a linear combination of these degenerate eigenmode pairs.

For these paired modes, we averaged the eigenvalue spectra and associated this average eigenvalue spectrum with the average of the corresponding eigenvalues.

#### Estimating dispersion relation

We used a heuristic procedure to identify the parameters of the wave equation. We used the eigenvalue spectra to identify a range of temporal stimulus frequencies (*ω)* associated with an eigenmode *k* and its corresponding Laplacian eigenvalue *η*_*k*_. This was done by fitting a locally-quadratic model to the most prominent peak in eigenvalue spectra (Figure 5C). To fit this model, we used all points between the “edges” of each peak, defined as the inflection points in the log-power spectrum adjacent to each local maximum. We used the peak of this quadratic fit as a proxy for a fundamental driving frequency *ω* associated with an eigenmode *k*. In cases when eigenvalue spectra contained multiple peaks, we selected a single peak satisfying the following constraints. First, we excluded the peaks that were 25% smaller than the maximal peak. Second, we assumed that Laplacian eigenvalues *η*_*k*_ and spatial frequency of evoked standing waves should monotonically increase with increasing flicker frequency *ω*, thus we discarded peaks that did not satisfy this constraint. We then combined data across animals and fitted dispersion relation derived from equation (1), thus mapping Laplacian eigenvalue *η*_*k*_ to the squared temporal frequency 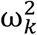 and estimating parameters *c* and *d* in equation (7).

## ACKNOWLEDGEMENTS

Rasa Gulbinaite has received funding from the European Union’s Horizon 2020 research and innovation programme under the Marie Skłodowska-Curie grant agreement No. 843379. Michael E. Rule is supported by a Leverhulme and Isaac Newton Trust fellowship ECF-2020-352. This work was also supported by the Natural Sciences and Engineering Research Council of Canada (No. 40352), Alberta Innovates, the Canadian Institute for Health Research (No. 390930), and Healthy Brains, Healthy Lives awarded to Majid H. Mohajerani.

## SUPPLEMENTARY MATERIALS

**Figure S1.**
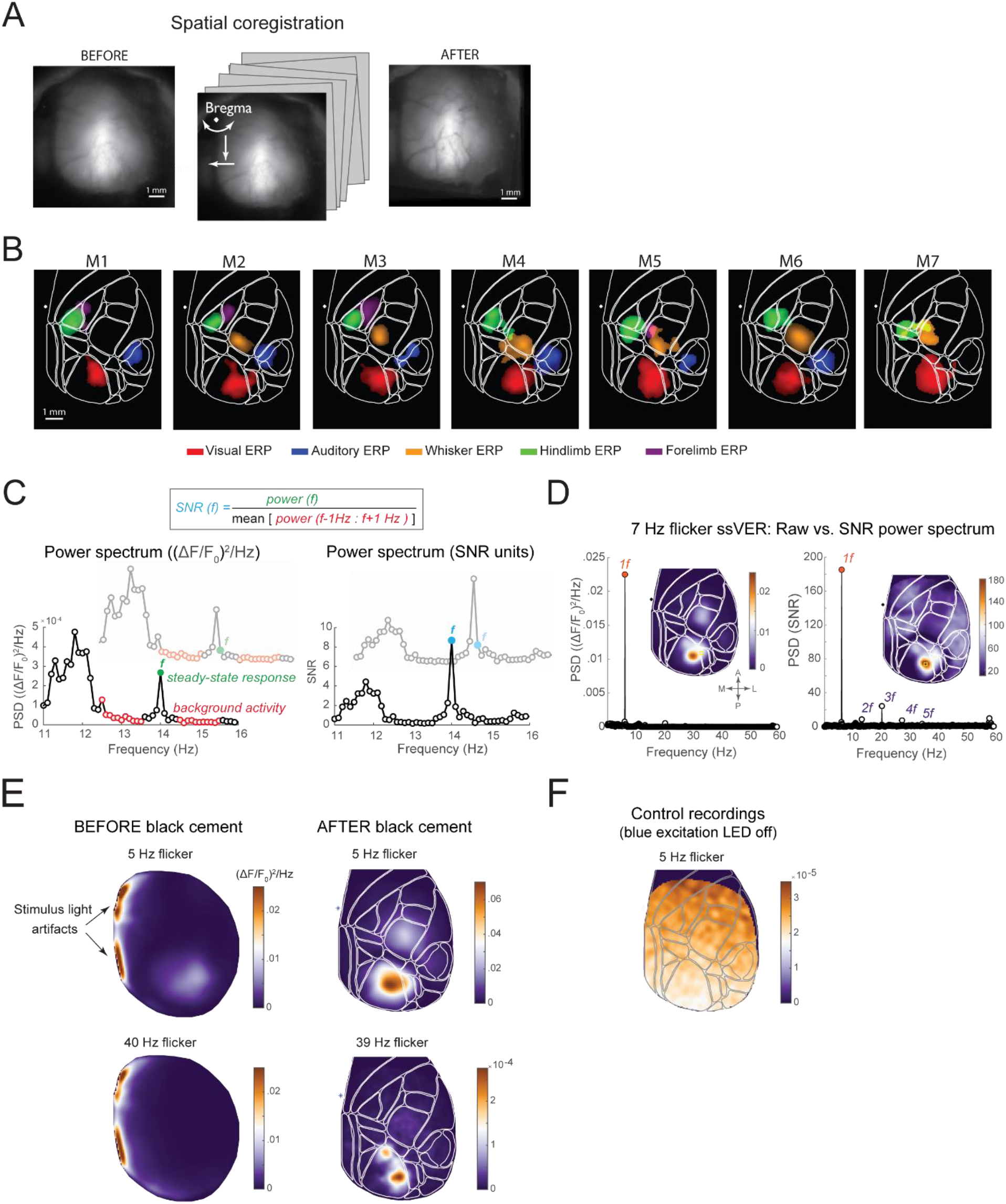
Data preprocessing, spatial co-registration across recordings and to Allen CCF (relates to Figures 1 and 2). (A)Reference images averaged across 5 recording sessions (performed across multiple days) before and after spatial co-registration. Blurry vasculature on the left indicates a slight variability of headfixation across recordings, which is corrected by rigid image transformation (translation and rotation) on the right (single animal data). (B)Functional maps from all animals representing an overlay of peak evoked activity for different sensory stimuli thresholded at 5 SD relative to the baseline (−1000 – -100 ms relative to the stimulus onset). Sensory stimulation recordings were co-registered to Allen CCF (white outlines) following the same procedure as ssVER recordings (see STAR Methods). High correspondence between anatomical areas in Allen CCF and activation peaks over primary sensory areas (VISp, AUDp, SSp-ll, SSp-up, SSp-bfd) shows the quality of alignment to Allen CCF. (C)High frequency flicker elicits smaller responses compared to low-frequency flicker due to 1/f power-law scaling (the steeper slope of background neural activity at slower frequencies). Therefore, the raw power spectra from each flicker condition were converted to SNR spectra to facilitate comparison of ssVERs evoked by different flicker conditions. Left: Illustration of the procedure for converting to SNR units. Each datapoint in the raw power spectrum is expressed as a ratio between the power at the frequency of interest and the average power at the neighboring frequencies (±1 Hz) of the FFT spectrum, excluding 0.5 Hz around the frequency of interest. Excluding immediate frequencies preserves the amplitude at the flicker frequency after conversion to SNR units, as ssVER signal is not a narrow-band sharp peak but also contains the side-bands. (D)Raw ((ΔF/F0)^2^/Hz units) vs. SNR power spectra from a single animal in response to 7 Hz flicker stimulation (averaged from 9 pixels around the spatial peak marked as yellow rectangles on the topographies). Note that SNR conversion reveals harmonic responses to flicker which are indistinguishable from background activity in the raw power spectrum. Topographies depict power at 7 Hz across the entire hemisphere. (E)Pilot recordings revealed that despite several measures we have taken to prevent stimulus LED light artifact in the imaging window (3D printed cone fitting the head-ring and black curtain attached to it), the left part of the imaging window was contaminated by the flickering stimulus light (arrows) positioned in the left hemifield. In these recordings, responses to low frequency 5 Hz flicker could be identified (top topography), while visual responses to 40 Hz flicker were not present (bottom topography). We therefore applied an additional layer of black cement over the frontal and lateral parts of the head-plate fixation and the skin. This procedure successfully eliminated light artifacts and revealed responses to high-frequency flicker (bottom topography). (F)5 Hz flicker control recordings performed at the end of each recording session during which the imaging window was not exposed to excitation blue LED. No neural fluorescence signal should be recorded under such conditions and only stimulus LED artifact should be visible in the imaging area. These control recordings showed that the power at the flicker frequency resulting from the LED visual stimulation was negligible and uniform across the imaging window (compare with the topographies on the left in E).

**Figure S2.**
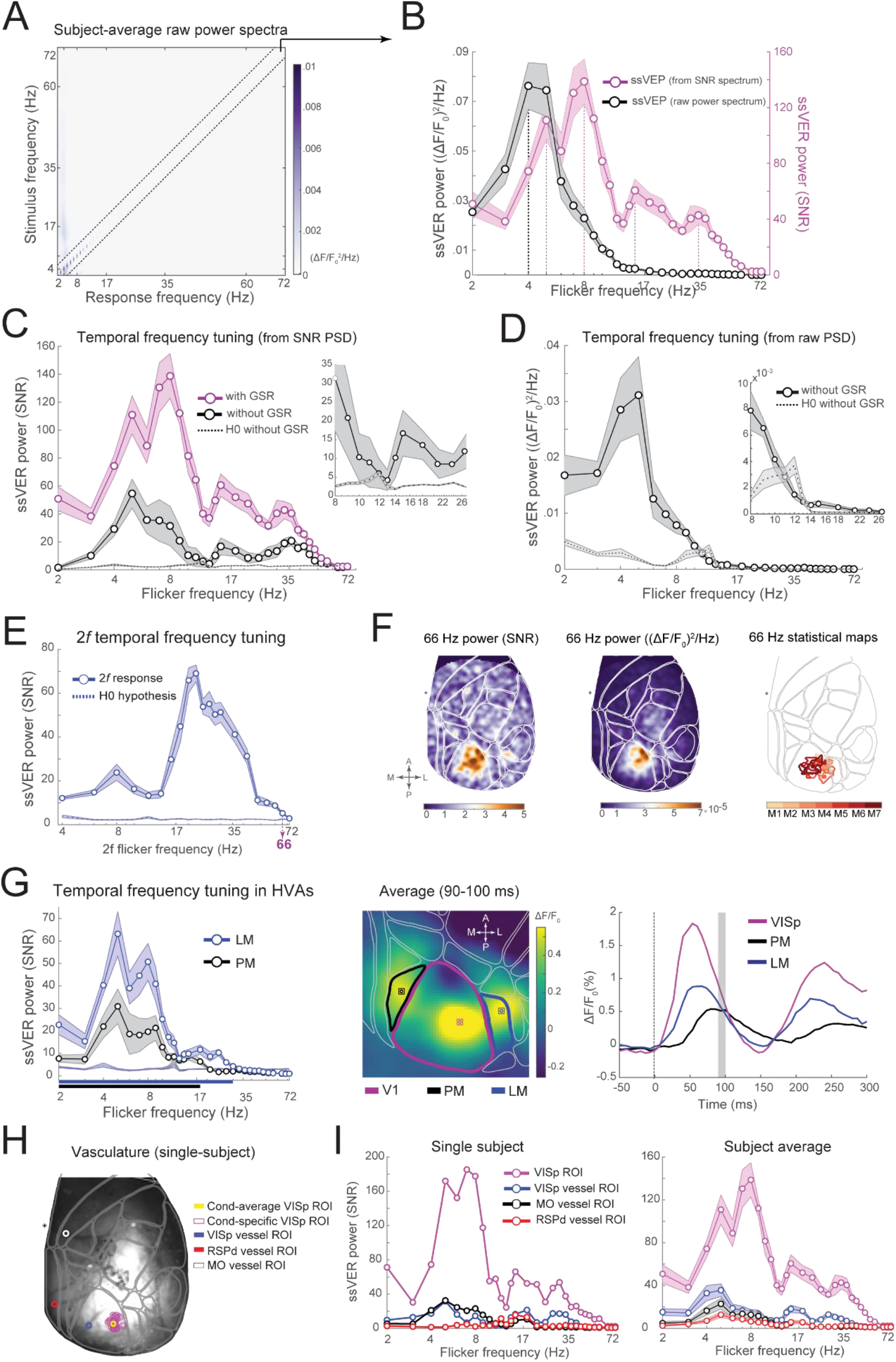
Temporal frequency tuning curves (relates to Figure 2). (A)Raw ((ΔF/F_0_^2^)/Hz units) subject-average power spectra (x-axis) in response to all flicker frequencies (y-axis). Note the apparent absence of responses to high-frequency flicker along the diagonal. (B)Temporal frequency tuning curves (data from the diagonal in A) derived from raw power spectra (black line) compared to SNR-converted power spectra (pink line). Note that only a single ∼5 Hz peak is visible in the temporal frequency tuning curves derived from the raw power spectra. (C) and (D) Effects of global signal regression (GSR) preprocessing step on temporal frequency tuning curves derived from the SNR-unit (C) and raw (D) power spectra. A strong heart-beat (8-14 Hz) signal is evident in the null-hypothesis curves (H0; dashed lines), which represent the background brain signal at flicker frequencies (computed from conditions when specific flicker frequency or its harmonics were not present). The difference between the pink and black lines in panel C indicates that GSR data preprocessing step increased the overall signal-to-noise ratio and successfully removed 8-14 Hz heartbeat signal revealing a resonance peak at ∼8 Hz. Insets: Zoomed-in plots of the power spectra showcasing the effects of GSR on 8-14 Hz activity. (E)Subject-average temporal frequency tuning curve for the 2^nd^ harmonic (2*f*) of the stimulation frequency shows that ssVER up to 66 Hz (33 Hz flicker stimulation) were statistically significant. The statistical threshold (95^th^ percentile of trials for which flicker frequency of interest was absent) is plotted as a dotted line on the bottom. For comparison, the 1^st^ harmonic responses (1*f*) were significant for flicker frequencies up to 56 Hz. (F)Left and middle: Exemplar ssVER topographies from a single subject at 66Hz, i.e. the 2^nd^ harmonic response to 33 Hz flicker (left: SNR, middle: raw power (ΔF/F_0_)^2^/Hz). Right: Spatial extent of statistically significant 2*f* responses. Data from different animals are plotted in different color contours. (G)Left: Temporal frequency tuning curves from two higher-order visual areas (HVAs): area PM (posteromedial) and LM (lateromedial). Solid lines on the x-axis indicate statistically significant responses up to 30 Hz in LM and up to 18 Hz in PM. In area LM, resonance peaks were more prominent and similar in frequency to that of V1. Middle: Area PM and LM ROIs were defined based on peak response times to single pulse stimuli. Right: Responses in PM and LM followed responses in V1 with a brief delay. (H)Vasculature image from a single subject illustrating ROIs positioned directly over the blood vessel in retrosplenial (red circle), motor (white circle), and visual cortices (blue circle from ROI over the blood vessel in VISp, purple circles indicate flicker frequency-specific ROIs and yellow one denotes flicker condition-average ROI). (I)Temporal frequency tuning curves from visual and non-visual ROIs positioned above blood vessels (single subject, same as in the main text figures) and subject average. Note that only visual ROIs show resonance peaks and neither motor nor retrosplenial ROI show resonance peaks around 8 Hz, 15 Hz, or 33 Hz. A characteristic 5 Hz peak is, however, present in all ROIs, and again indicates a potentially non-visual nature of this rhythmic activity, which likely is a breathing rhythm as discussed in the main text and Figure S3.

**Figure S3.**
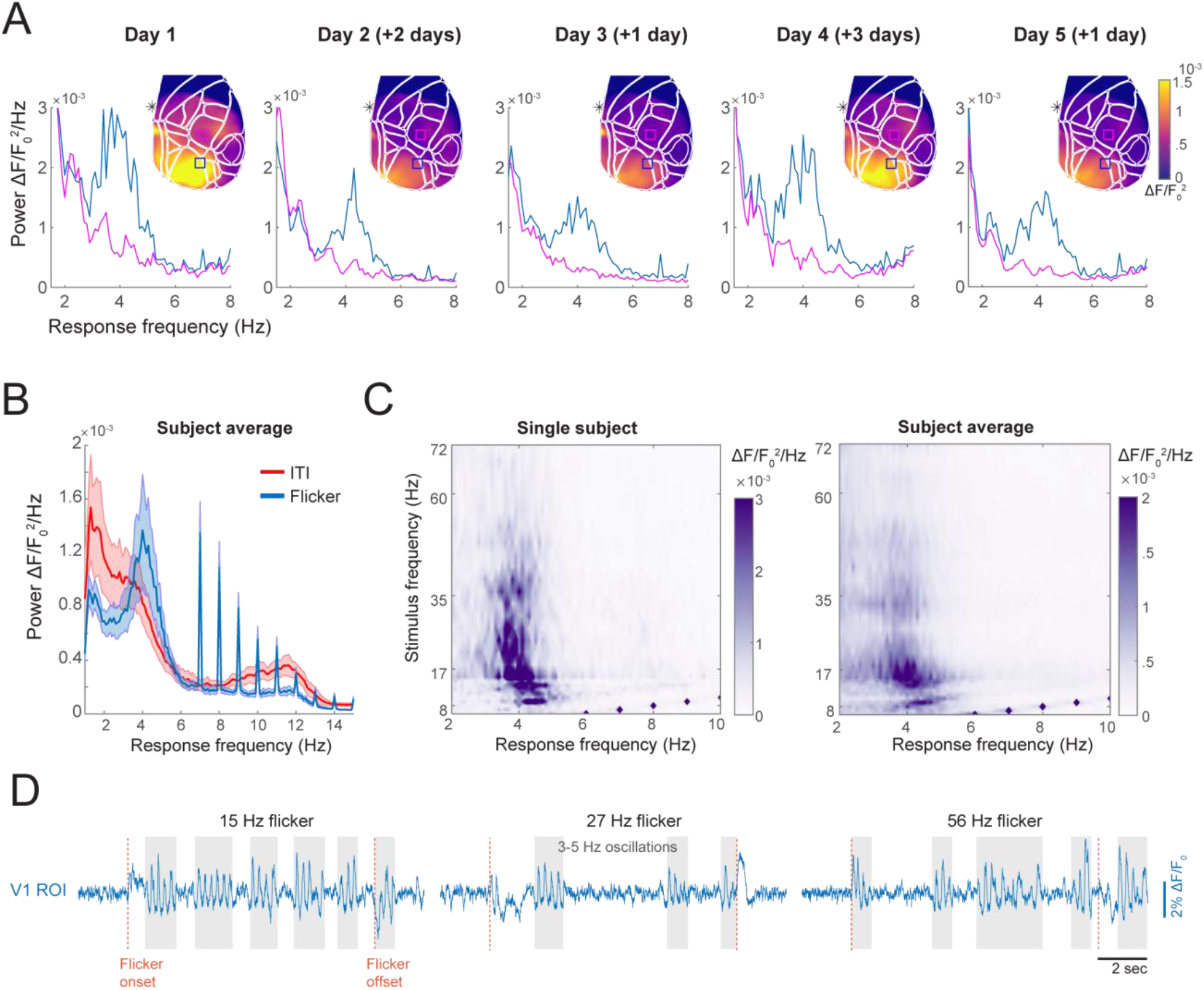
Origins of 5 Hz resonance peak in temporal frequency tuning curves (relates to Figure 2). (A)Power spectra from V1 ROI (blue trace) and S1 ROI (magenta trace) and topography of 3-5 Hz activity during >7 Hz flicker trials from a single animal across recording days. The 3-5 Hz activity was more prominent over visual areas (power differences between blue and magenta traces). To make sure that 3-5 Hz activity in the visual cortex was not a result of global signal regression (GSR) pre-processing step, the data plotted here does not include GSR preprocessing step. The numbers in the brackets indicate the number of days relative to the previous recording and thus the recency of exposure to the flickering stimulus. (B)3-5 Hz rhythm was more prominent during the stimulus than during inter-trial interval (ITI). (C)The power of 3-5 Hz rhythm varied as a function of frequency, with lower flicker frequencies eliciting stronger 3-5 Hz power as compared to the higher flicker frequencies. (D)Illustration of V1 ROI time series during three different flicker frequency trials (single animal data).

**Figure S4.**
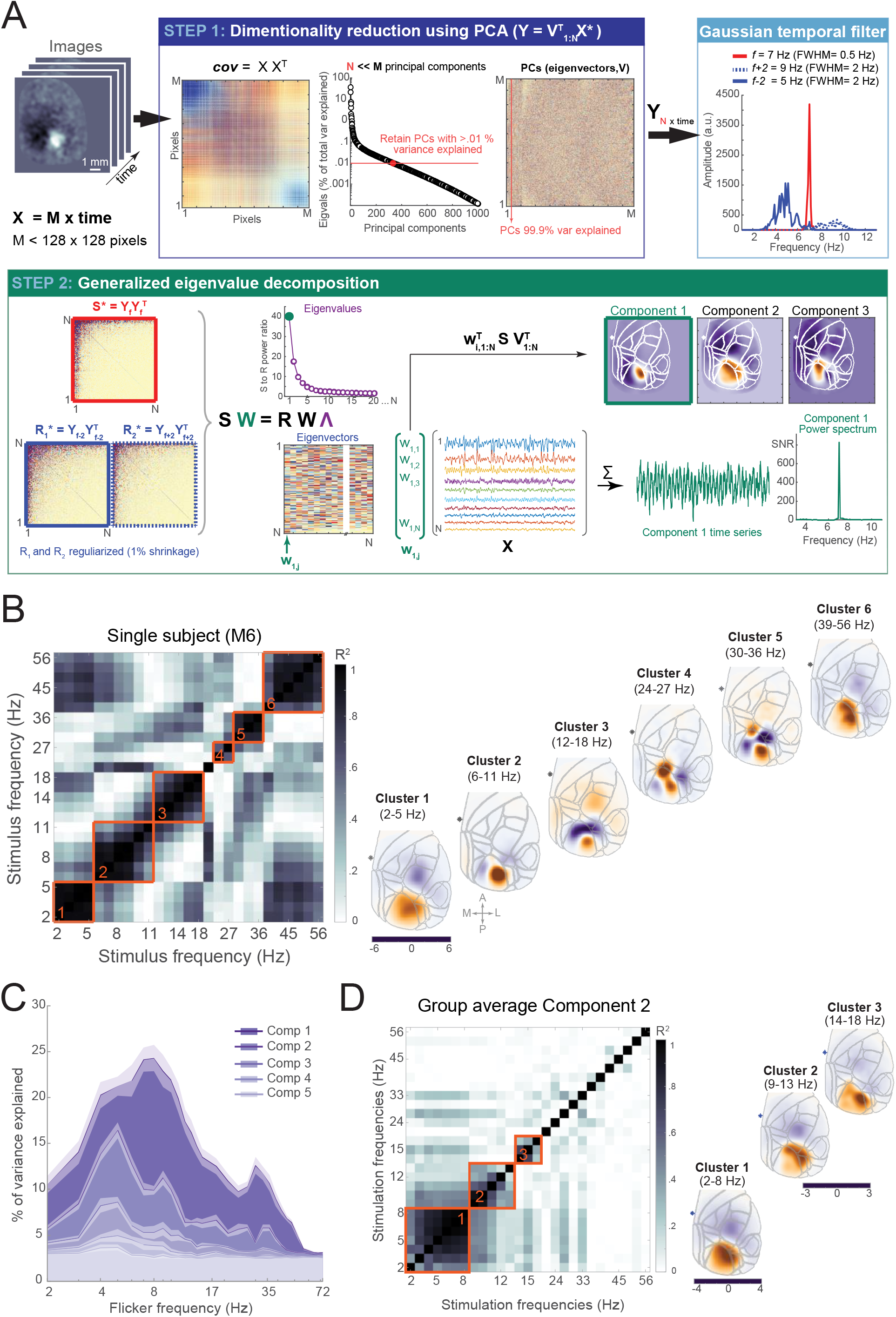
Spatiotemporal response pattern analysis (relates to Figure 3). (A)Illustration of two-stage generalized eigenvalue decomposition (GED) used to isolate spatiotemporal response patterns characteristic to each flicker frequency. In Step 1, a 1000-10000 ms time window of single-trial baseline-corrected fluorescence time series **X** (M x time, where M is the number of pixels containing brain data) were used to compute single-trial covariance matrices (M x M), which were averaged and submitted to PCA. Eigenvectors (principal components) associated with eigenvalues that explain more than 0.01% variance were used to reconstruct principal component time series **Y** (N x time, where N is the number of retained PCs and N << M, e.g. 364 vs. 8798). Single-trial time series **Y** were then narrow bandpass-filtered around the stimulation frequency *f* (FWHM = 0.5 Hz) and nearby frequencies *f ± 2 Hz* (FWHM = 2 Hz). In Step 2, 1000-10000 ms time window of temporally filtered data was used to compute signal covariance matrix **S** (N x N) and two reference covariance matrices **R** (N x N) that were averaged. The spatial filters that optimally separate covariance matrices **S** and **R** were computed using GED, where **W** is a matrix of eigenvectors, and **Λ** is a diagonal matrix of eigenvalues. Eigenvector w_i,j_ (marked in blue in matrix **W**) associated with the highest eigenvalue λ (marked in green in diagonal of **Λ** plot) was used to reconstruct cortical maps and component time series. (B)Left: Same as Figure 3G in the main text but for a single animal. Pairwise squared correlation matrix across spatiotemporal patterns (derived using the analysis procedure described in A) generated in response to different flicker frequencies. The hierarchical clustering procedure identified 6 distinct clusters of ssVER topographies for this animal, as depicted on the right. Note the similarity of cluster 4 and 5, which, for other animals, was identified as a single cluster. (C)Although the 1^st^ GED component captures the most variance, GED decomposes the signal into as many components as there were PCs included in the analysis (on average 377 components). Variance explained by the first 5 components is plotted for all flicker conditions. Although the 1^st^ component was chosen because it explains most variance and best separates the S from R matrix, some variance was also explained by the 2^nd^ component. (D)Contrary to the 1^st^ GED component, 2^nd^ component spatiotemporal response patterns were spatially homogenous.

**Figure S5.**
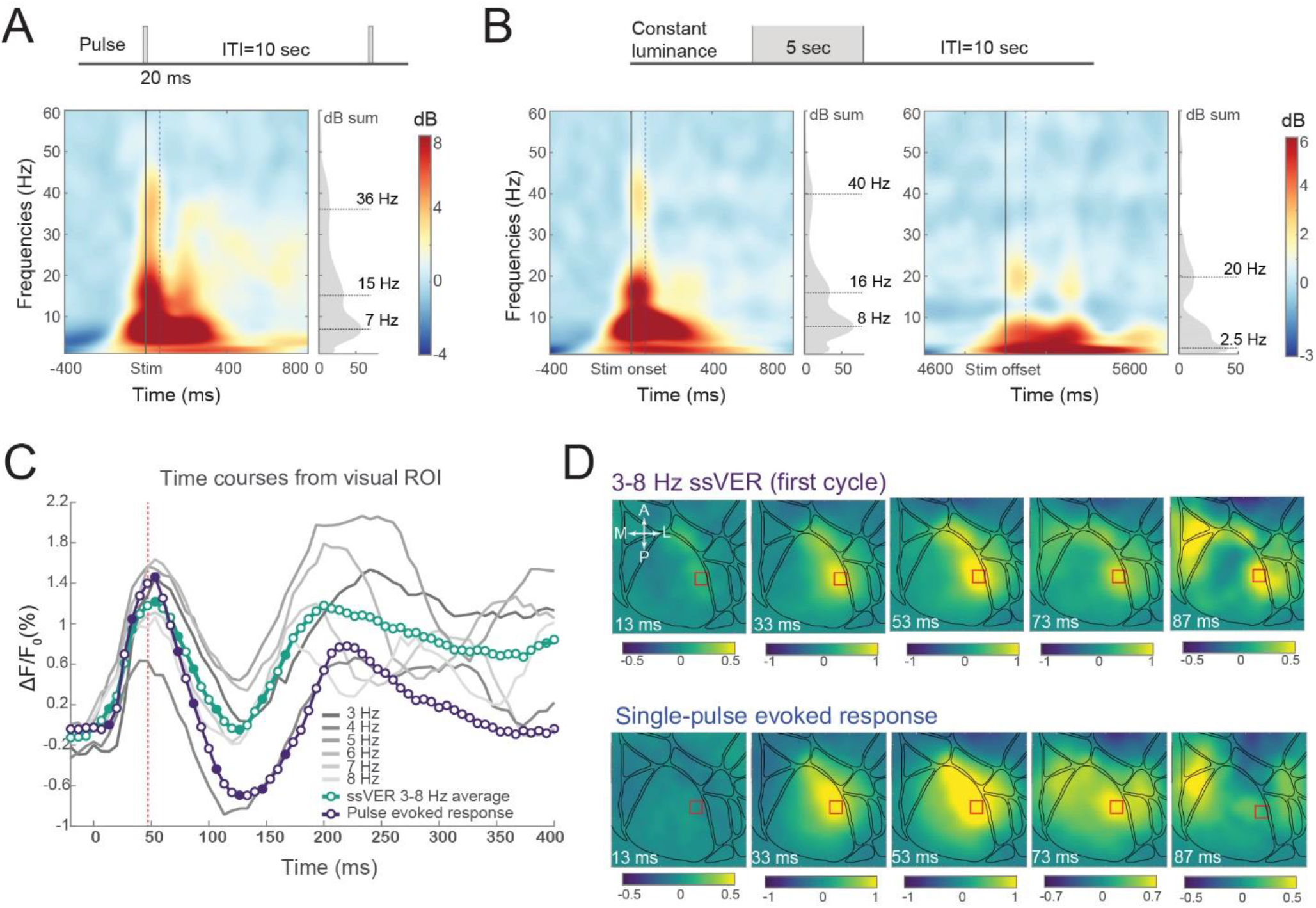
Similarities between ssVER and evoked responses to single-pulse and constant luminance stimulation (relates to Figure 4). (A) and (B) Baseline-correct time-frequency representations of responses to pulse and constant luminance stimuli (subject-average data). Data plotted from V1 ROI (9 pixels around the spatial peak evoked response). The left and the right graph in B represents response to luminance increase (trial onset) or decrease (trial offset) respectively. Spectra in the insets represent power values summed over time window marked with dashed lines (either 0-70 or 0-100 ms in the rightmost graph). Time-frequency decomposition was performed separately for each trial using wavelet transformation, then the trials were averaged and baseline-corrected relative to -500 – -100 ms (where 0 is the stimulus onset). Wavelet transform parameters used in the analysis: 80 linearly spaced frequencies ranging from 1 to 60 Hz in 1 Hz steps, number of wavelet cycles ranged from 3 to 17 and was also linearly spaced. Following pulse and onset of continuous luminance stimuli there was an increase in theta (peak M_pulse_ = 6.97 Hz; M_ON-OFF_ = 7.72 Hz), beta (M_pulse_ = 15.19 Hz; M_ON-OFF_ = 15.94 Hz, and gamma power (M_pulse_ = 36.1 Hz; M_ON-OFF_ = 39.84 Hz). (C) Similarities between the first cycle of 3-8 Hz ssVER and evoked responses to single-pulse stimulation. Time courses are from the ROI marked in panel D, which was defined based on the spatial peak of response in each paradigm separately. (D) Time-lapse images of stimulus-evoked activity during the first cycle of 3-8 Hz sine-wave stimulation (top row) and evoked responses to single-pulse stimulation (bottom row) from the solidly colored timepoints in panel C.

**Figure S6.**
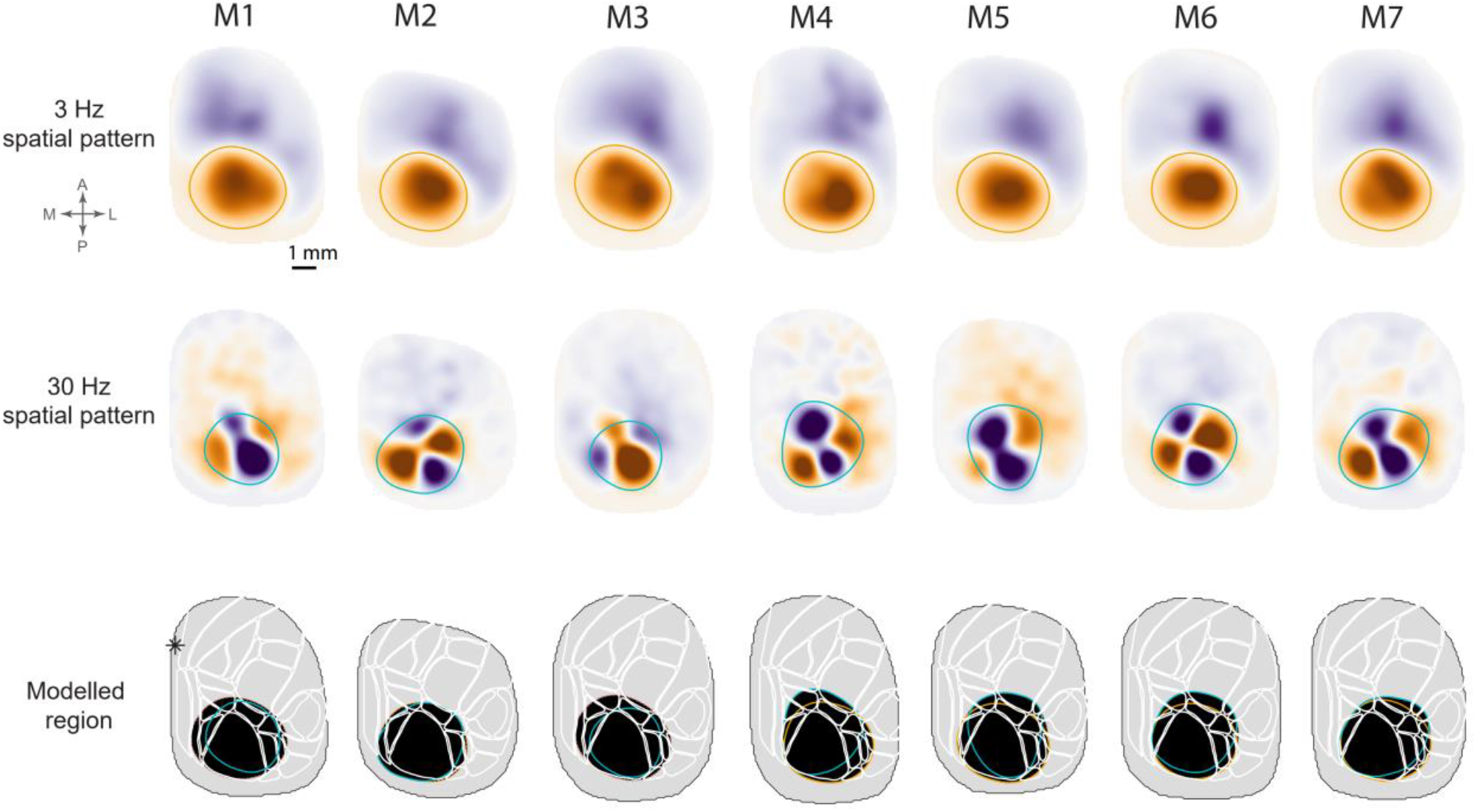
Region of cortex used for modeling spatial eigenmodes (relates to Figure 5). Given the clearly defined near zero amplitude contours, spatiotemporal response patterns identified using two-stage GED from two flicker conditions (3 Hz and 30 Hz) were used to define a region of posterior cortex subsequently used for modeling geometric eigenmodes. The union of regions (orange for 3 Hz condition and green for 30 Hz condition) defined from both conditions was used (black area in the bottom plots).

## Notes

https://osf.io/u37ar/

### Competing Interest Statement

The authors have declared no competing interest.

### Summary of Updates

A few changes in the Results section.

